# Early-life Microbiome-Derived HCA Enhances Type 3 Immunity via FXR- dependent m^6^A RNA methylation

**DOI:** 10.1101/2024.07.04.602137

**Authors:** Zhipeng Yang, Zhiyuan Lin, Yaojie You, Mei Zhang, Ning Gao, Xinru Wang, Jian Peng, Hongkui Wei

## Abstract

The gut microbiome plays a crucial role in the development of intestinal immunity during early life, but the underlying mechanisms remain largely unknown. Here, we found oxygen consumption in neonatal rats by *S. boulardii* accelerated the colonization of the microbiome and the development of type 3 immunity, which protected against *S. typhimurium*. Microbiome maturation increased the abundance of microbiome-encoded bile salt hydrolase (BSH) genes and elevated the levels of Hyocholic acid (HCA). HCA promotes the development of γδT and type 3 innate lymphoid cell (ILC3) by sustaining the stability of *Rorc* mRNA via the FXR-WTAP- N6-methyl-adenosine (m^6^A) axis. *L. reuteri*, a bacterium encoding BSH, was enriched by oxygen consumption in the intestine and promoted intestinal type 3 immunity. However, inhibition of BSH blocks the *L. reuteri-*induced development of intestinal type 3 immunity. These results reveal the role of microbiome-derived HCA in the regulation of intestinal type 3 immunity during early life.

## Introduction

Approximately 4% of children under the age of 5 die worldwide. Despite the gradual decline in the mortality rate in recent years, it remains at a high level in some developed countries, particularly in Asia and Africa [1]. Infections significantly contribute to early postnatal mortality in neonates. This is largely attributed to the immature immune system in neonates, which makes them more vulnerable to infections than adults [2]. Early life is a crucial stage for the development of the neonatal immune system. Numerous immune cells amplify in the early stage, including T cells [3] and innate lymphoid cells (ILCs) [4], playing a rural role in the pathogen infection. Type 3 immunity is characterized by an immune response with a distinct profile, including the expression of genes encoding IL-17A and IL-22, and the key transcription factor retinoic acid-related orphan receptor (RORγt), mainly consisting of ILC3, γδT cells, and Th17 [5]. Type 3 immunity serves a vital role in defending against microbial infection and maintaining intestinal epithelial homeostasis [6].

Early life is a critical period for mammals to be colonized with the microbiome, which profoundly impacts the development of intestinal immune function. Early life exposure to antibiotics leads to an immature microbiome and type 3 immune system [7]. Moreover, germ-free models showed that the function of the type 3 immune system depends on the intestinal microbiome [3, 8]. Short-chain fatty acids (SCFAs) and tryptophan metabolism support expansion and cytokines secretion of ILC3 and T cells [9]. The intestinal microbiome also influences host epigenetic modification [10]. Microbial transfer to mice pups restored histone modifications in the ileum [11]. In addition, the presence of microbes inhibits the host m^6^A modification, as demonstrated by antibiotic and germ-free mouse models [12]. Mechanically, microbial metabolite folate regulates the m^6^A modification level by providing SAM, and butyrate participates in the tricarboxylic acid cycle for the demethylation of RNA [13]. Various studies have reported that m^6^A is a major post-transcriptional regulator of immune cells [14]. Whereas, whether the microbiome contributes to the regulation of m^6^A modification in intestinal immune cells in early life remains unknown.

Previous research supports the idea that oxygen is consumed by pioneer facultative anaerobic bacteria in the aerobic gut in early life, which then provides a niche for obligate anaerobes to colonize [15]. Here, we demonstrate that intestinal luminal oxygen consumption in the early life of rats facilitates the colonization of the microbiome, which leads to a better ability to cope with *Salmonella typhimurium* (*S. typhimurium*) via type 3 immune response. HCA enhances the stability of *Rorc* mRNA via WTAP-mediated m^6^A via inhibition of FXR and GW4064 offset the regulatory effect of HCA on the development of type 3 immunity. *L. reuteri* enriched the abundance of HCA and enhanced type 3 immune response depending on BSH. Collectively, these data suggest a mechanism by which early-life microbiome influences the host type 3 immunity, with HCA produced by *Lactobacillus* affecting the stability of *Rorc* mRNA through m^6^A modification in an FXR-dependent manner.

## Result

### Early intervention with *S. boulardii* accelerated oxygen consumption in the intestinal lumen and facilitated the maturation of the intestinal microbiome

To investigate the effects of intestinal oxygen consumption on microbial colonization in early life, we performed early intervention with the anaerobic microorganism, *Saccharomyces boulardii* (*S. boulardii*), in neonatal rats and assessed changes in the gut oxygen status and gut microbial colonization. The findings revealed a gradual decline in intestinal oxygen levels as the organism matured. Early intervention with *S. boulardii* significantly reduced oxygen levels at 3 days and trended to decreased oxygen levels at 8 days (**Figure 1A**). The hypoxic environment strongly stabilizes HIF-1α and transactivates HIF target genes such as *Bnip3l* in epithelial cells [16]. In the current study, early intervention with *S. boulardii* reduces mRNA expression of HIF-1α and target genes, including *Pfkb3*, *Defb1*, *Muc3*, *Bnip3l*, *Slc2a1* and *Tff3* at 3, 8, 14 days in the ileum, indicating a relatively hypoxic environment in the intestinal lumen (**Supplementary** Figure 2A-C). For alpha-diversity of the microbiome, early intervention with *S. boulardii* significantly increased the Chao1 index at 3 days after birth and significantly decreased the Chao1 index at 8 days (**Supplementary** Figure 3A). At the beta-diversity, the SB group showed a significant difference compared to the Con group, especially after 5 days (**Supplementary** Figure 3B-G).

**Figure 1.**
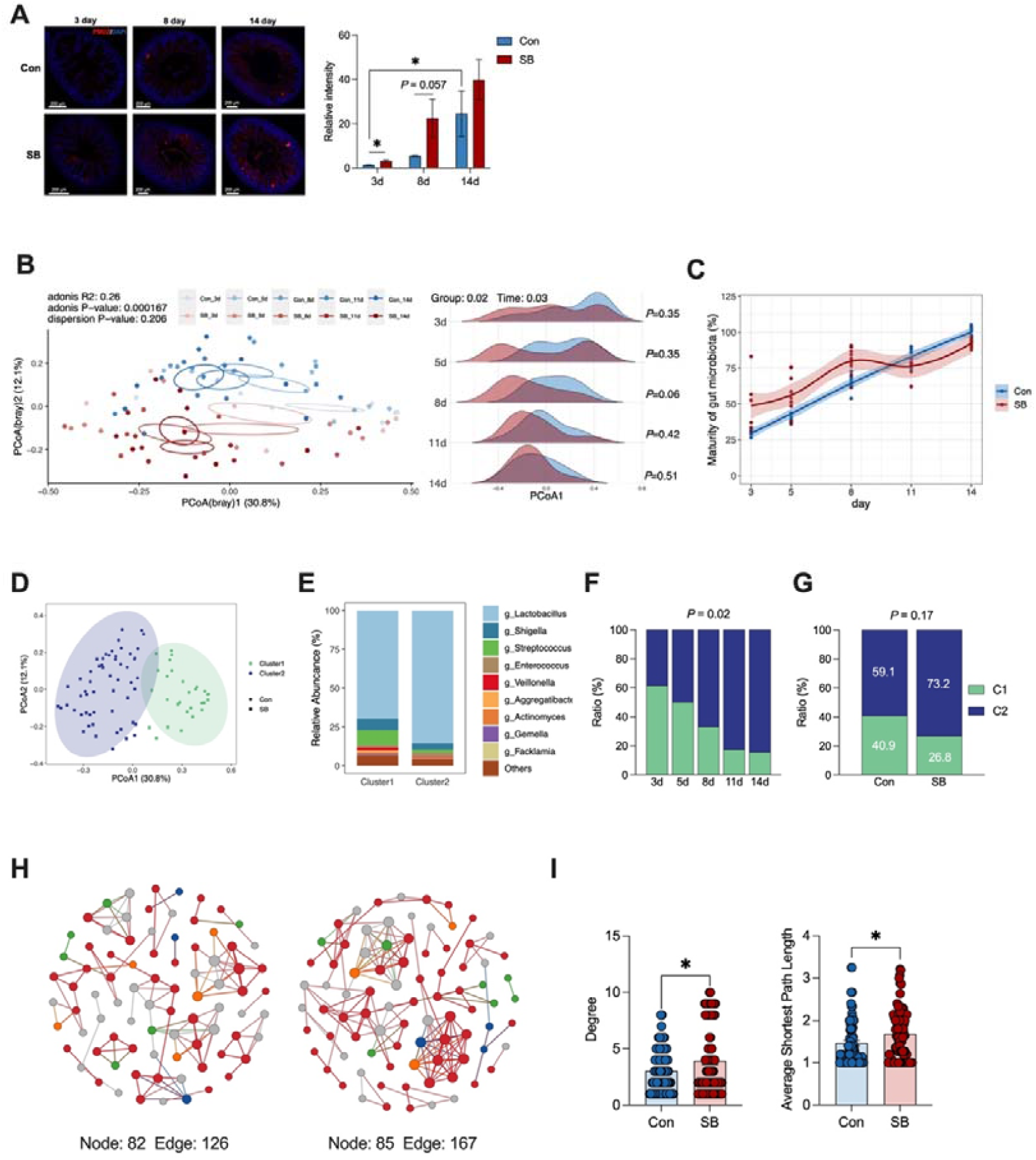
*S. boulardii*-induced oxygen consumption facilitates the maturation of the microbiome in the intestine of neonatal rats (A) Representative images of Immunofluorescent staining for Hypoxyprobe-1(red) and nuclear (blue) in ileum tissue and quantification of staining intensities. n = 3-4. (Scale bars: 200 μm). (B) PCoA plot of rats after PBS and *S. boulardii* treatment at days 3, 5, 8, 11, and 14 after birth on the Bray-Curtis. The density plot shows a comparison of the PcoA1 of the Con groups and SB group at days 3, 5, 8, 11, and 14 after birth. (C) Intestinal microbiome maturity curve of rats after PBS and *S. boulardii* treatment by random forest. (D) Two clusters were identified by an unsupervised clustering algorithm. Cluster 2 was identified as a mature type, and Cluster 1 was identified as an immature type. (E) Intestinal microbiome composition at the genus level of Cluster 2 and Cluster 1. (F-G) Proportions of the two clusters at the five sampling times (F) and comparison of the proportions of the two clusters of rats after PBS and *S. boulardii* treatment (G). The significance of dissimilarity was calculated by chi-squared test. (H) Network analysis of the intestinal microbiome of rats after PBS and *S. boulardii* treatment. (I) Comparison of network degree and average shortest path length. Data represent mean ± SEM. \**P*[<[0.05. The difference between rats after PBS and *S. boulardii* treatment was analyzed by two-tailed unpaired Student’s t-test, except for (C) (D) (F) (G).

To evaluate the effect of *S. boulardii*-induced oxygen consumption on microbiome maturation, we utilized permutational multivariate analysis of variance (PERMANOVA) on Bray-Curtis dissimilarities among samples. Both individual age and *S. boulardii* had a significant impact on the microbiome composition (**Figure 1B**). Interestingly, a transition in the microbiome was observed as time points increased in the PCoA1 axis, which was accelerated in neonates receiving *S. boulardii*, particularly at 8 days (**Figure 1B**). Furthermore, a random forest was conducted to match the microbiome maturity curve, revealing that *S. boulardii* treatment accelerated microbiome development (**Figure 1C**). Next, an unsupervised clustering algorithm was used, and the neonatal intestine microbiome was classified into two clusters, which also corresponded to the maturation trajectory observed in the PCoA plot (**Figure 1D**). At the genus level, we observed that the *g_Lactobacillus* was enriched in Cluster 2 (mature type), while *g_Streptococcus* was enriched in Cluster 1 (immature type) (**Figure 1E**). The ratio of mature microbiome (cluster 2) gradually increased during growth (**Figure 1F**), and a higher proportion of mature microbiome (cluster 2) was observed in the SB group compared to the Con group (**Figure 1G**). Early life microbial maturation often involves an increase in community diversity and interaction network complexity [17]. To further assess community ecological parameters, subsequent analysis by co-occurrence network analysis to compare the complexity and stability between the Con and SB groups. The SB group had more nodes and edges compared to the Con group (**Figure 1H**), and at the node level, the degree and average shortest path length of the SB group were significantly higher than those of the Con group (**Figure 1I**). Collectively, these analyses suggest that the oxygen consumption by *S. boulardii* accelerates the assembly of the microbiome towards a more mature and stable microbiome.

### *S. boulardii-*induced oxygen consumption facilitated the maturation of type 3 immune cells

The early establishment of the intestinal microbiome in life is crucial for the maturation of immune cells. Therefore, we detected the transcriptional levels of three transcription factors *Tbx21*, *Ggta3*, and *Rorc*, which respectively direct the maturation of type 1, 2, and 3 immune cells [18]. Among these, only the mRNA level of *Rorc* was elevated after early intervention with *S. boulardii* (**Supplementary** Figure 4A). In addition, the mean fluorescence intensity (MFI) of RORγt in the intestinal lamina propria lymphocytes (LPLs) of the SB group also increased (**Figure 2A**). Next, we examined the proportion of type 3 immune cells, Th17, γδT, and ILC3 in LPLs. The proportion of γδT cells and ILC3 (**Figure 2B, C**), but not Th17 (**Supplementary** Figure 4B) were significantly elevated after early intervention with *S. boulardii*. Type 3 immune cytokines IL-17A and IL-22 in γδT cells and ILC3 were also improved after *S. boulardii* early intervention (**Supplementary** Figure 4C-F). The proportion of T cells, B cells, Macrophages, CD4^+^T cells, and CD8^+^T cells was not changed (**Supplementary** Figure 4G**, H**).

**Figure 2.**
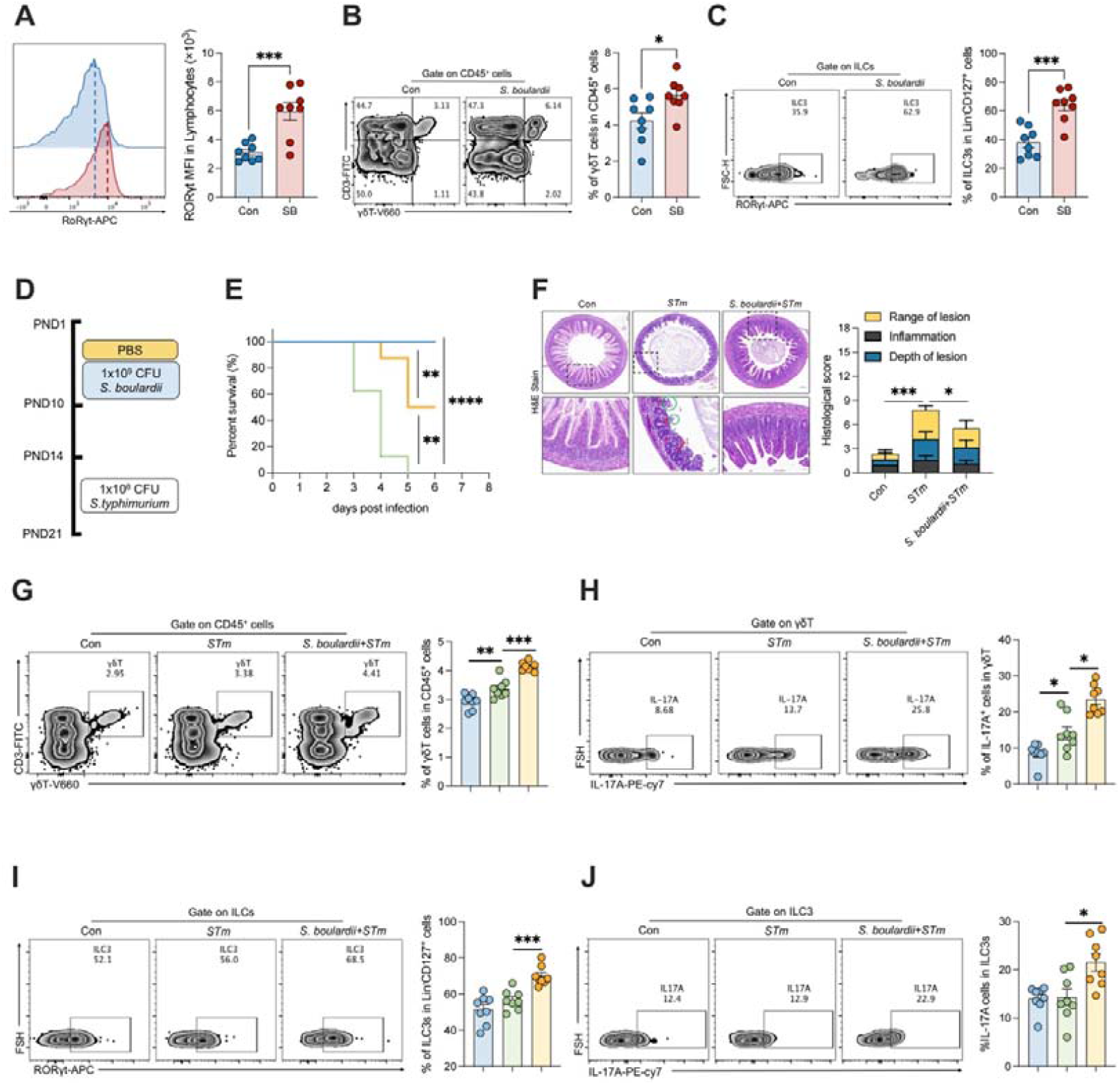
*S. boulardii*-induced oxygen consumption facilitated the maturation of type 3 immune cells and enhanced resistance to *S. typhimurium* infection (A) Representative FACS plots and MFI of RORγt expression in CD45^+^ cells of small intestine LPLs of rats. n = 8. (B-C) Representative FACS plots and % of γδT and ILC3 in small intestine LPLs of rats. n = 8. (D) Workflow of *S. boulardii* treatment and *S. typhimurium* infection in neonatal rats (E) Survival curve of the Con, SB, and SB + *STm* groups. The significance of dissimilarity was calculated by the Log-rank (Mantel-Cox) test. n = 6. (F) Representative H&E images showing histological scores in Con, SB, and SB + *STm* groups. n = 5 - 6. (G-H) Representative FACS plots and % of γδT and IL-17A^+^γδT in small intestine LPLs of rats. n = 8. (I-J) Representative FACS plots and % of ILC3 and IL-17A^+^ILC3 in small intestine LPLs of rats. n = 8. Data represent mean ± SEM. \**P*[<[0.05; \*\**P*[<[0.01; \*\*\**P*[<[0.001 The difference between the two groups was analyzed by two-tailed unpaired Student’s t- test, , except for (E).

### *S. boulardii*-induced oxygen consumption confers protection from *S. typhimurium infection*

Next, we investigated whether oxygen consumption by *S. boulardii* protects against *salmonella* infection. All control rats succumbed to *S. typhimurium* infection within 5 days post-infection (dpi), while approximately 50% of rats supplemented with *S. boulardii* survived (**Figure 3E**). Early intervention with *S. boulardii* reduced weight loss and diarrhea in neonatal rats after *S. typhimurium* infection (**Supplementary** Figure 5A**, B**). *S. typhimurium* colonization in the small intestine (jejunum and ileum) was reduced by *S. boulardii* intervention but not in the large intestine (cecum and colon), and the liver at 1 dpi (**Supplementary** Figure 5C). However, *S. boulardii* significantly reduced *S. typhimurium* colonization in the ileum and liver at 3 dpi (**Supplementary** Figure 5D). *S. typhimurium* infection induced extensive epithelial damage and inflammatory cell infiltration in neonatal rats, which was alleviated by *S. boulardii* intervention (**Figure 2F**).

**Figure 3.**
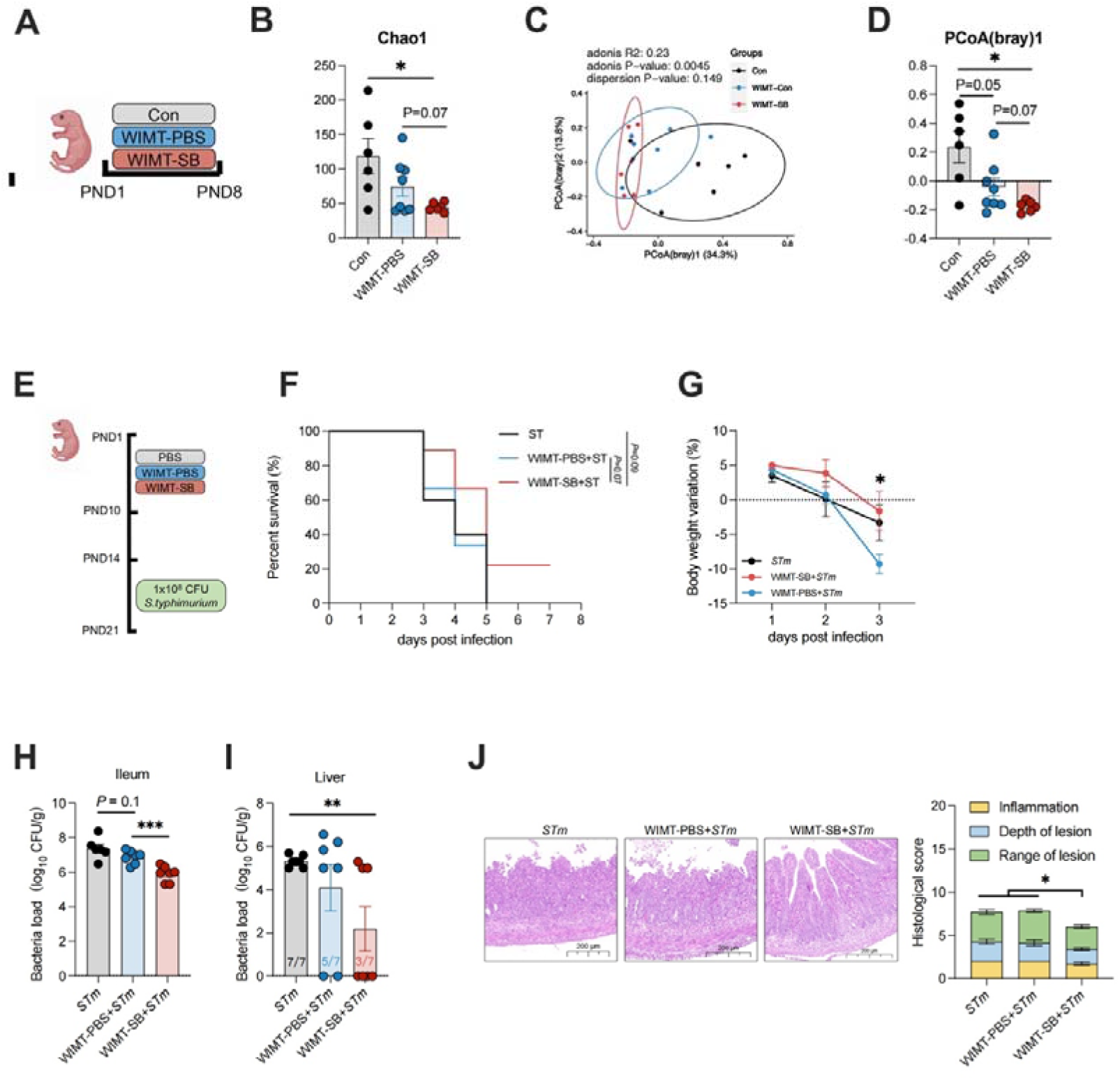
Microbial transfer in the SB group accelerates microbial maturation and enhances resistance to *S. typhimurium* infection (A) Workflow of WIMT treatment in neonatal rats (B) The α-diversity Chao1 index and β-diversity of ileum content microbiome (C-D) PCoA (principal coordinate analysis) plot of rats from three groups on the Bray-Curtis. The significance of dissimilarity was calculated by adonis and dispersion. (E) Workflow of WIMT treatment and *STm* infection in neonatal rats (F) Survival curve of the *STm*, WIMT-PBS+*STm*, and WIMT-SB+*STm* groups. The significance of dissimilarity was calculated by the Log-rank (Mantel-Cox) test. n = 9. (G) Bodyweight variation of the *STm*, WIMT-PBS+*STm*, and WIMT-SB+*STm* groups. n = 9. (H-I) *STm* burden in the Ileum and Liver of the *STm*, WIMT-PBS+*STm*, and WIMT- SB+*STm* groups. n = 7. (J) Representative H&E images showing histological scores in *STm*, WIMT- PBS+*STm*, and WIMT-SB+*STm* groups. n = 7. Data represent mean ± SEM. \**P*[<[0.05; \*\**P*[<[0.01; \*\*\**P*[<[0.001. The difference between the two groups was analyzed by two-tailed unpaired Student’s t- test, except for (C) (F).

The previous study demonstrated that *S. boulardii* directly restricts *S. typhimurium* expansion in the intestine [19]. Therefore, we detected the *S. boulardii* load in the intestinal content and found that *S. boulardii* was not present in the intestinal content 4 days after stopping *S. boulardii* intervention (**Supplementary** Figure 5E), suggesting effect on *S. typhimurium* colonization was not due to the direct function of *S. boulardii*. Then, we examined the type 3 immune response in neonatal rats after *S. typhimurium* infection. *Rorc* and *Il17a* mRNA levels were upregulated in the SB group, but there was no significant difference in *Il22* (**Supplementary** Figure 5F). Similarly, the protein level of IL-17A, but not IL-22 was upregulated in the SB group (**Supplementary** Figure 5G). Moreover, *Il17a* mRNA levels correlated with the bacterial load in the ileum (**Supplementary** Figure 5H). Additionally, the type 3 immune response, including the percentage of γδ T cells and ILC3 and their ability to secrete IL-17A, was enhanced by *S. boulardii* treatment (**Figure 2G-J**). The results showed that *S. boulardii* reduces *S. typhimurium* lode, alleviates weight loss and diarrhea, and increases the proportion of IL-17A secreting-γδT cells and ILC3.

### Microbial transfer in the SB group accelerates microbial maturation and enhances resistance to *S. typhimurium* infection

To investigate whether the oxygen consumption-induced maturation of intestinal type 3 immunity depends on the microbiome, we conducted whole intestinal microbial transplantation (WIMT) (**Figure 3A**), which is more conducive to donor small intestinal microbiome colonization [20]. Chao 1 index in the WIMT-SB group was lower than in the Con group (**Figure 3B**) but was similar to that in the WIMT-PBS group. The β-diversity result showed that the microbial composition was different in the three groups (**Figure 3C, D**).

Next, we explored the effect of WIMT on *S. typhimurium* infection (**Figure 3E**). All neonatal rats from the Con and WIMT-PBS groups succumbed to *S. typhimurium* infection within 5 days post-infection, while approximately 20% of neonatal rats from the WIMT-SB group survived (**Figure 3F**). Similarly, WIMT-SB significantly alleviated body weight loss and *S. typhimurium* loading in neonatal rats post-infection compared to WIMT-PBS (**Figure 3G-I**). Additionally, WIMT-SB group rats exhibited lower histological scores than the *STm* and WIMT-PBS groups (**Figure 3J**). Overall, we found that WIMT from the SB group accelerates microbial maturation and enhances resistance to *S. typhimurium* infection in neonatal rats, indicating that the oxygen consumption-induced maturation of intestinal type 3 immunity depends on the microbiome.

### Gut microbial maturation facilitates type 3 cell function through BAs metabolism

To explore the potential functional capabilities of the mature microbiome, KEGG function prediction using PICRUSt2 analysis was conducted. We found secondary bile acid biosynthesis was the most enriched pathway and had the highest fold change in the top 10 KEGG pathways (**Figure 4A**). The level of secondary bile acid biosynthesis gradually increased with growth and was significantly higher at 14 days than at 3 days. Moreover, the level of secondary bile acid biosynthesis in the SB group was higher than that in the Con group at 8 and 11 days (**Figure 4B**). Therefore, the profiles of BAs in the ileum of rats were detected by LC-MS/MS. A total of 63 BAs were detected with clear separation in BA profile between SB and Con groups by OPLS-DA analysis (**Figure 4C**). Overall, the absolute level of total BAs and primary BAs decreased in the SB group (**Figure 4D**), while relative levels of primary BAs decreased significantly, and secondary BAs increased significantly in SB group (**Figure 4E**). Among the 63 BAs measured, 7 differential BAs were identified as significantly different (Fold change > 2, *P* < 0.05). These included elevated 3β-HDCA and HCA, and decreased GCDCA-3S, TLCA-3S, TCA, TLCA, and TCDCA (**Figure 4F**). Focusing on elevated HCA and 3β-HDCA, absolute HCA level was significantly higher than 3β-HDCA (**Figure 4G**). Moreover, 3β-HDCA was not detected in two samples, thus we hypothesized that HCA may contribute to the crosstalk between the microbiome and the immune maturation of the host.

**Figure 4.**
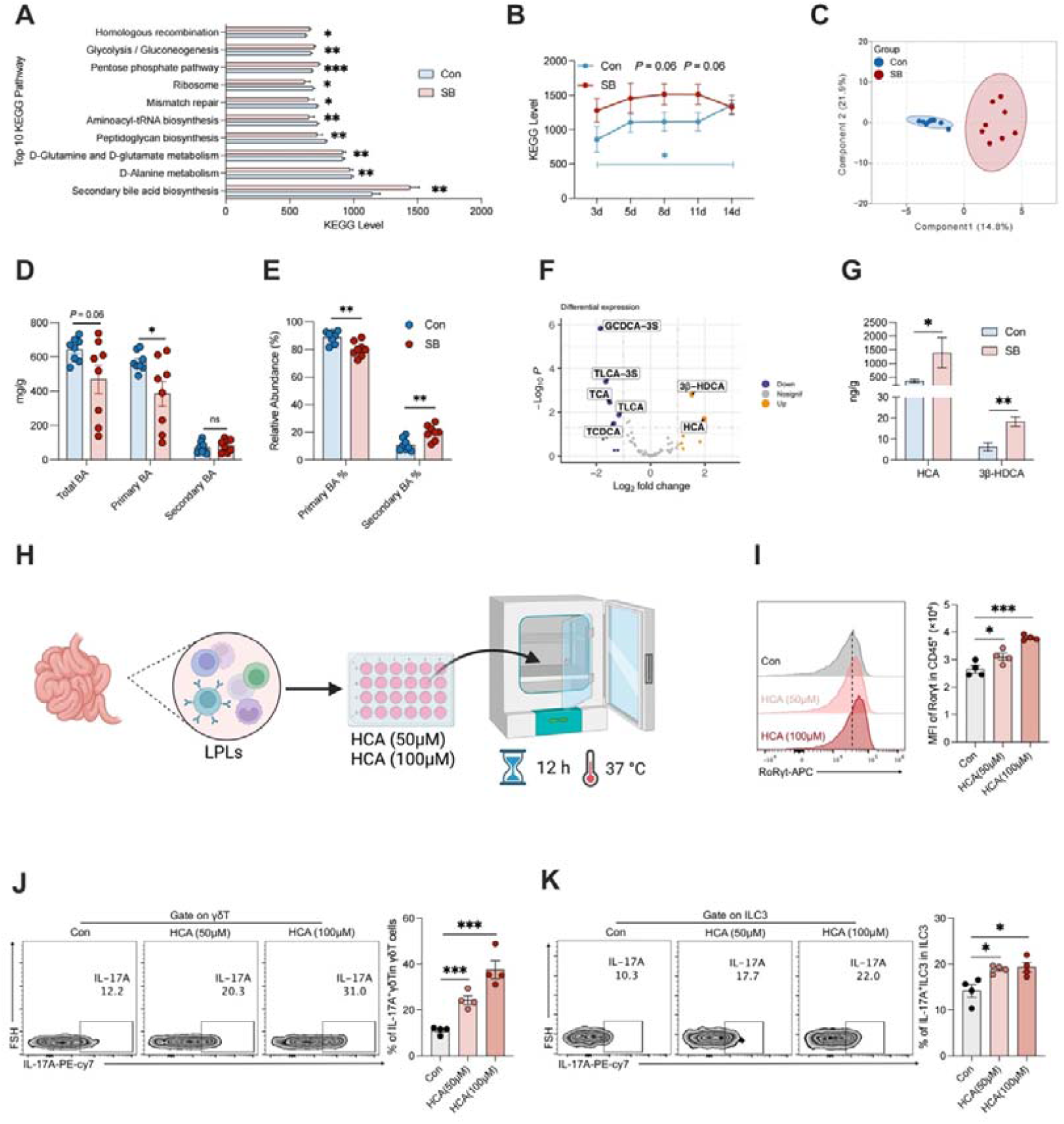
Gut microbial maturation facilitates type 3 cell function through BAs metabolism (A) Top 10 predicted microbial functions based on PICRUSt2 between rats after PBS and *S. boulardii* treatment. n = 44 and 41. (B) The secondary BA biosynthesis level between rats after PBS and *S. boulardii* treatment at the five-time points. n = 6 - 10 (C) OPLS-DA score plot of the Ileum content BA profiles between rats after PBS and *S. boulardii* treatment at 8 days. n = 8. (D) The absolute abundance of primary BA, secondary BA, and total BA. n = 8. (E) The relative abundance of secondary BA and primary BA. n = 8. (F) Volcano plot analysis of different BA with a threshold of FC > 2 or FC < 0.5, and *P*-value < 0.05. n = 8. (G) The absolute abundance of enriched BA in the SB group, HCA, and 3β-HDCA. n = 8. (H) Workflow of HCA treatment on LPLs from neonatal rats HCA in vitro. (I) Representative FACS plots and MFI of RORγt expression in CD45^+^ cells of small intestine LPLs of rats after HCA treatment in vitro. n = 4. (J and K) Representative FACS plots and % of ILC3 and IL-17A^+^ILC3 in small intestine LPLs of rats after HCA treatment in vitro. n = 4. Data represent mean ± SEM. \**P*[<[0.05; \*\**P*[<[0.01; \*\*\**P*[<[0.001. The difference between the two groups was analyzed by two-tailed unpaired Student’s t- test, except for (B) (C) (F)

To verify this, we isolated LPLs from the small intestine of rats and incubated them ex vivo with HCA (**Figure 4H**). The MFI of RORγt was increased by HCA treatment (**Figure 4I**). Furthermore, the ability of IL-17A secreting of ILC3 and γδT was enhanced after HCA treatment (**Figure 4J, K**). Together, these data suggest that secondary BAs biosynthesis is enriched in the intestinal microbiome after *S. boulardii* intervention, and HCA enhances the function of type 3 immune γδT cells and ILC3.

### HCA enhances the function of type 3 immune cells via inhibition of FXR

Previous research reported that HCA is an antagonist of farnesoid X receptor (FXR) [21], and blocking FXR activation promotes IL-17A production in ILC3 [22]. Therefore, we hypothesize HCA enhances the function of type 3 immune cells via FXR. We found HCA treatment with LPLs increased the FXR target gene (*Fgf19* and *Shp*) expression (**Figure 5A**). To fully elucidate the interaction between HCA and FXR, a molecular docking study was conducted. The docking results showed HCA could bind well to FXR through visible hydrogen bonds and strong electrostatic interactions, with a low binding energy of -7.991 kcal/mol. The hydrogen bonding between the residue SER-332 and HCA played an important role in the observed interaction (**Figure 5B**).

**Figure 5.**
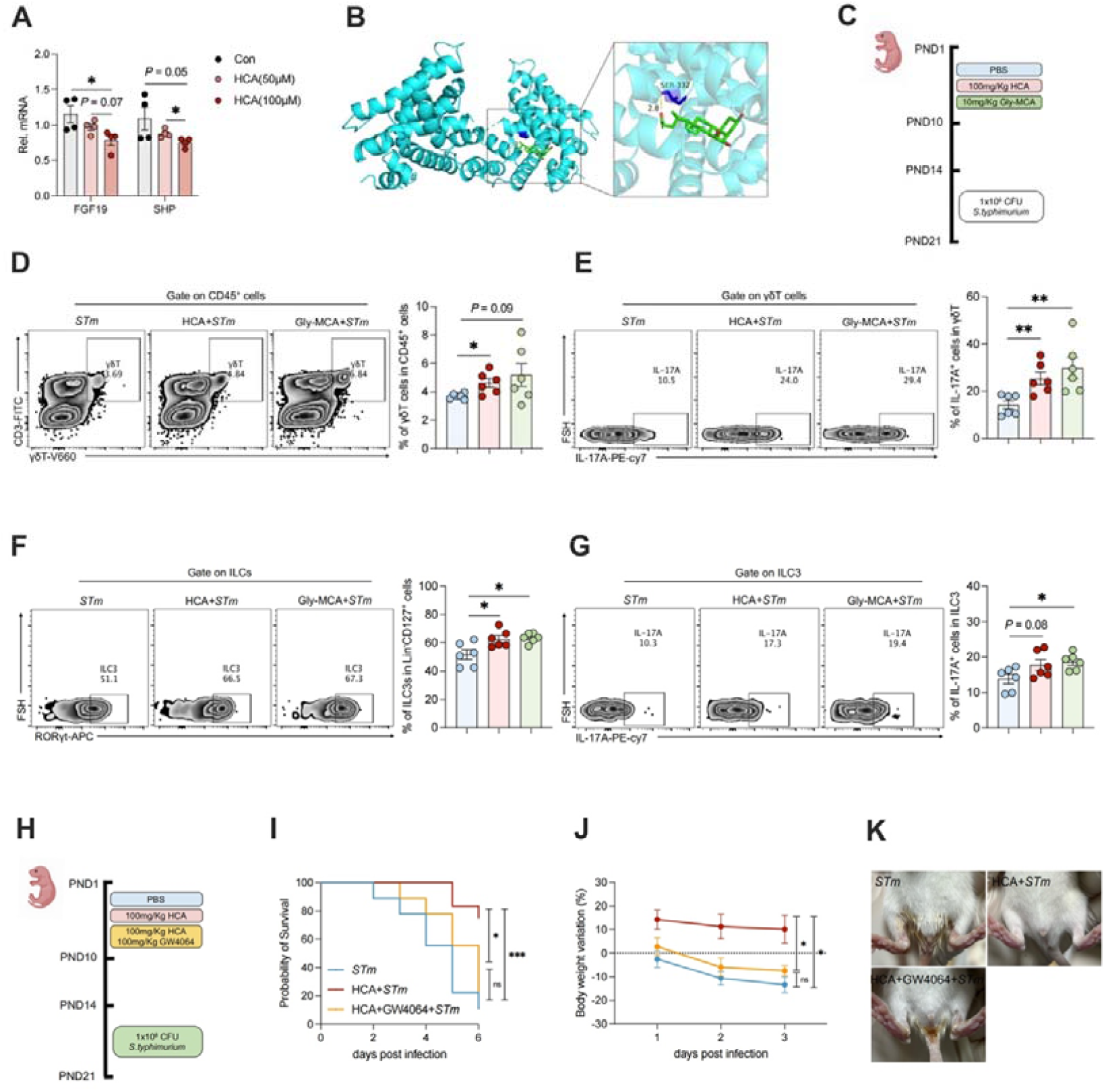
HCA enhances the function of type 3 immune cells via inhibition of FXR (A) RT-qPCR results of mRNAs encoded by the FXR target gene *Fgf19 and Shp* in LPLs after HCA treatment in vitro. n = 4. (B) Molecular docking shows HCA combines at the FXR SER-332. (C) Workflow of HCA and Gly-MCA treatment and *STm* infection on neonatal rats (D-E) Representative FACS plots and % of γδT and IL-17A^+^γδT in small intestine LPLs of *STm*, HCA + *STm*, Gly-MCA + *STm* groups. n = 6. (F-G) Representative FACS plots and % of ILC3 and IL-17A^+^ILC3 in small intestine LPLs of *STm*, HCA + *STm*, Gly-MCA + *STm* groups. n = 6. (H) Workflow of HCA and GW4064 treatment and *STm* infection on neonatal rats (I) Survival curve of the *STm*, HCA + *STm*, HCA + GW4064 + *STm* groups. The significance of dissimilarity was calculated by the Log-rank (Mantel-Cox) test. n = 9- 12. (J) Bodyweight variation of the *STm*, HCA + *STm*, HCA + GW4064 + *STm* groups. The significance of dissimilarity was calculated by t-test. n = 9-12. (K) Representative diarrhea images of *STm*, HCA + *STm*, HCA + GW4064 + *STm* groups. Data represent mean ± SEM. \**P*[<[0.05; \*\**P*[<[0.01; \*\*\**P*[<[0.001; ns, no significance. The difference between the two groups was analyzed by two-tailed unpaired Student’s t-test, except for (I).

Therefore, we orally administered HCA and intestine-selective FXR inhibitor glycine- β-muricholic acid (Gly-MCA) [23] to neonatal rats (**Figure 5C**). Both HCA and Gly- MCA alleviated the mortality, weight loss, and diarrhea induced by *S. typhimurium* infection (**Supplementary** Figure 6A-C). Further, the type 3 immune response, including the percentage of γδ T cell and ILC3 and the ability of IL-17A secretion were enhanced by HCA and Gly-MCA (**Figure 5D-G**). We then treated rats with GW4064, a general FXR agonist, in combination with HCA to test the mediated effect of FXR (**Figure 5H**). GW4064 substantially counteracted the protective function of HCA against *S. typhimurium* including mortality, loss of body weight, and diarrhea (**Figure 5I-K**). Collectively, our data suggest that HCA protects against *S. typhimurium* through type 3 immune response via inhibition of FXR.

### HCA enhances the mRNA stability of *RORC* by inhibition of FXR-induced WTAP transcription

Given that *Rorc* is a vital factor for the development of type 3 immunity, and *S. boulardii*-induced oxygen consumption promotes *Rorc* expression in LPLs, we investigated the effect of HCA or Gly-MCA treatment on *Rorc* expression in LPLs. Increased *Rorc* expression was observed in LPLs of rats treated with HCA or Gly-MCA (**Supplementary** Figure 7A). Although FXR is a nuclear receptor, previous studies have shown that FXR does not transcriptionally regulate *Rorc* [22]. Therefore, we speculate that other epigenetics may contribute to the connection between FXR and *Rorc*. m^6^A is one of the most common modifications of RNA molecules in mammalian cells and broadly affects immune responses [24]. Furthermore, the modification of m^6^A in the intestine can be regulated by the microbiome [12]. To verify the importance of m^6^A modification in the development of microbiome and immune cells, we first detected the change in m^6^A modification of LPLs. The global m^6^A level in LPLs mRNAs was lower in 14-day-old rats than in 3-day-old rats after birth (**Supplementary** Figure 7B), suggesting that m^6^A modification plays a vital role in the development of immune cells. We also found that *S. boulardii* decreased the m^6^A level in the LPLs (**Supplementary** Figure 7C). We hypothesized that FXR may affect m^6^A modifications through transcriptional regulation. Therefore, we detected the mRNA levels of m^6^A-related enzymes, and the results showed that the three m^6^A readers, *Mettl3*, *Mettl14*, and *Wtap*, were downregulated after *S. boulardii* treatment (**Supplementary** Figure 7D). In addition, the treatment of HCA and GW4064 decreased the m^6^A level and the *Wtap* level but not the *Mettl3* and *Mettl14* levels of LPLs (**Figure 6B**). We also predicted the specific binding of FXR in the promoters of *Wtap* by the Jasper database.

**Figure 6.**
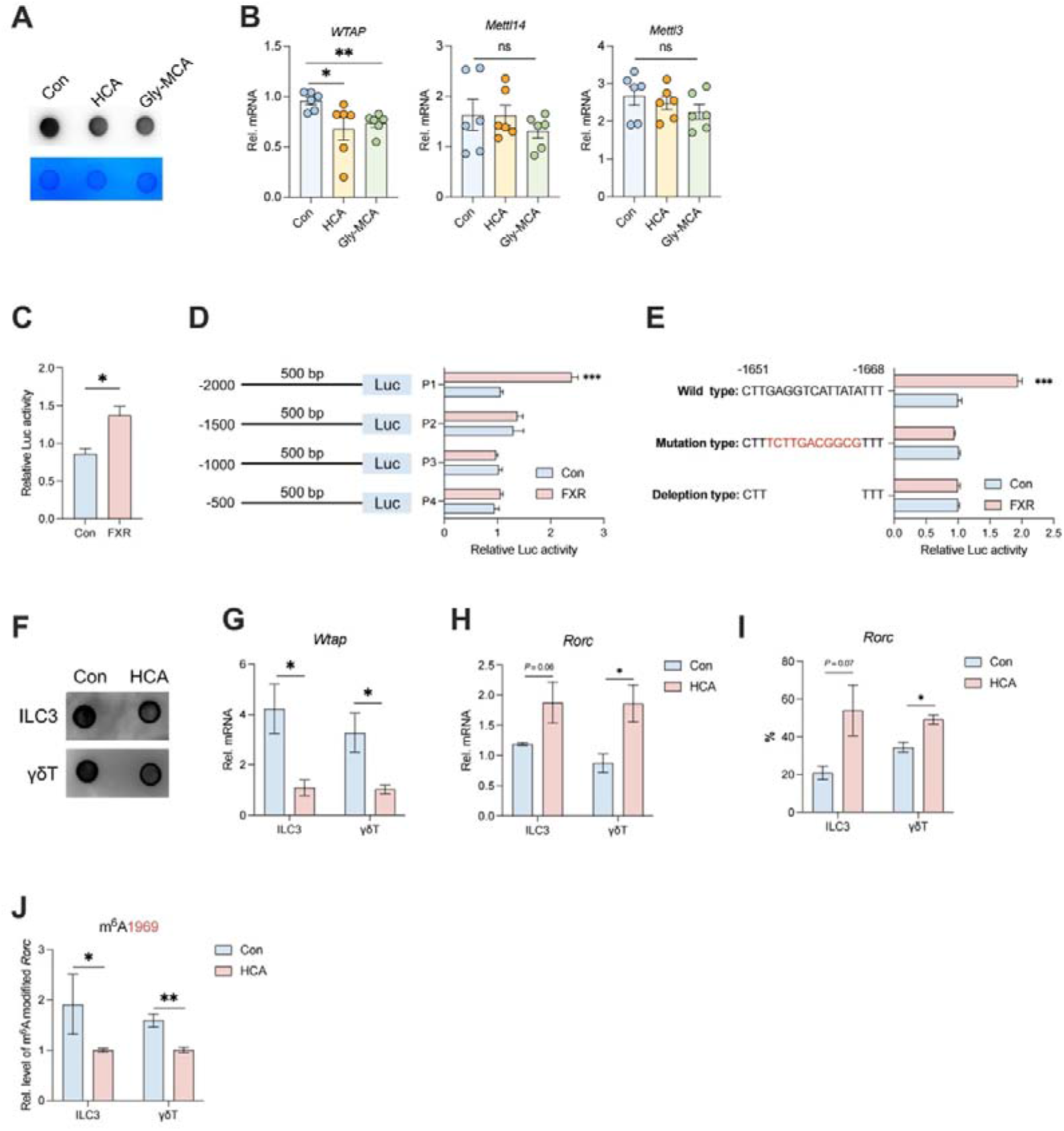
HCA enhances the mRNA stability of *RORC* (A) Dot blot to detect the m^6^A level of mRNA isolated from total RNA of LPLs isolated from rats from Con, HCA, and Gly-MCA groups. (B) RT-qPCR results of mRNAs encoded by the m^6^A methylases gene *Wtap, Mettl3, and Mettl14* in LPLs isolated from rats from Con, HCA, and Gly-MCA groups. (C) The *Wtap* promoter reporter was co-transfected with CMV-FXR or empty vector (pReceiver-M35) into 293T cells, and the promoter activity was determined as the ratio of firefly/Renilla luciferase activities. The dual-luciferase activity was measured after 24 hours of transfection. (D) Relative promoter activity of *Wtap* promoter reporter (P1-P4). (E) Relative promoter activity of the WT of *Wtap* promoter and the *Wtap* mutation and *Wtap* deletion constructs. (F) Dot blot to detect the m^6^A level of mRNA isolated from total RNA of rat intestinal γδT and ILC3 with or without HCA treatment. (G-H) RT-qPCR results of mRNAs encoded *Wtap* and *Rorc* genes in rat intestinal γδT and ILC3 with or without HCA treatment. (I) RNA stability of *Rorc*. RT-qPCR results of mRNAs encoded *Rorc* genes in rat intestinal γδT and ILC3 with or without HCA treatment compared 0 hour to 5 hours. (J) m^6^A level at 1969 of mRNA of *Rorc* detected by Epi-SELECTTM m^6^A site identification kit. n = 3. Data represent mean ± SEM. \**P*[<[0.05; \*\**P*[<[0.01; \*\*\**P*[<[0.001. The difference between the two groups was analyzed by two-tailed unpaired Student’s t- test.

To confirm whether FXR transcriptionally regulates *Wtap*, a luciferase reporter plasmid of the human *Wtap* promoter and an FXR cDNA plasmid were constructed for a dual-luciferase assay. FXR overexpression increased the luciferase activity of the *Wtap* reporter (**Figure 6C**). To identify the region responsible for FXR-mediated transactivation, we created constructs containing fragments of the human *Wtap* promoter (P1-P4), each of them co-transfected into 293T cells with FXR overexpression plasmids. As shown in **Figure 6D**, FXR overexpression significantly enhanced the activity of the P1 promoter (-2000 bp to -1500 bp) in the reporter assay, but not P2-P4. When this binding site was mutated or deleted in the *Wtap* promoter, FXR-mediated transactivation of promoter activity was abolished (**Figure 6E**). Thus, these results suggest FXR plays a crucial role in WTAP transactivation.

To determine whether the m^6^A pathway contributes to HCA regulating ILC3 and γδT, we sorted the ILC3 and γδT cells and treated them with HCA. As expected, HCA reduced the global m^6^A levels and the expression of *Wtap* in ILC3 and γδT cells (**Figure 6F**, **G**). Additionally, HCA increased the expression of *Rorc* in LC3 and γδT cells (**Figure 6H**). RNA decay assay showed that HCA treatment slowed the *Rorc* transcript decline (**Figure 6I**). The 12 potential m^6^A sites in the *Rorc* mRNA was predicted by using SRAMP and the decreased m^6^A modification levels of *Rorc* in HCA-treated LC3 and γδT cells was verified using Epi-SELECTTM (**Figure 6J)**. Collectively, these data indicate that FXR transcriptionally regulates m^6^A methyltransferase WTAP and affects the *Rorc* mRNA m^6^A modification and its stability.

### *L. reuteri* enriched HCA in neonatal rats

Next, we aimed to identify the bacteria contributing to the enrichment of HCA in neonatal rats. Our analysis revealed a positive correlation between the abundance of *Lactobacillus* and secondary BA biosynthesis (**Supplementary** Figure 8A). Moreover, the abundance of *Lactobacillus* was also correlated positively with HCA (**Supplementary** Figure 8B). The first step in the metabolism of secondary BAs is hydrolysis by bile salt hydrolase (BSH). Given that *Lactobacillus* is one of the BSH- expressing bacteria, we compared the relative abundance of *Bsh* in the microbiome between the Con and SB groups. Compared to Con, the SB group showed a tendency for increased *Bsh* relative abundance (**Supplementary** Figure 8C). All enriched ASVs sequences in the SB groups were annotated as *Lactobacillus* (**Supplementary** Figure 8D). Additionally, *L. reuteri* and *B. animals* were the enriched species in the WIMT-SB (**Supplementary** Figure 8E). Among them, only *L. reuteri* was enriched in both SB and WIMT-SB rats (**Supplementary** Figure 8F). Then, *L. reuteri* enrichment in the SB group at 8 days was validated by using qPCR assay (**Supplementary** Figure 8G).

To test the effect of *L. reuteri* on BAs metabolism, we treated neonatal rats with *L. reuteri* or control media and analyzed the BAs profiles in the intestine at 8 days of life (**Figure 7A**). OPLS-DA analysis demonstrated a clear separation in the BAs profile between the two groups (**Figure 7B**). The absolute level of total BAs and primary BAs decreased (**Figure 7C**). Furthermore, the relative levels of primary BAs had a trend to decrease, and the relative levels of secondary BAs had a trend to increase after *L. reuteri* treatment (**Figure 7D**). Among secondary BAs, HCA levels was increased after *L. reuteri* treatment (**Figure 7E; Supplementary** Figure 8H). Collectively, our findings suggest that *L. reuteri* modulated the BA profiles and enriched HCA in intestine of neonatal rats.

**Figure 7.**
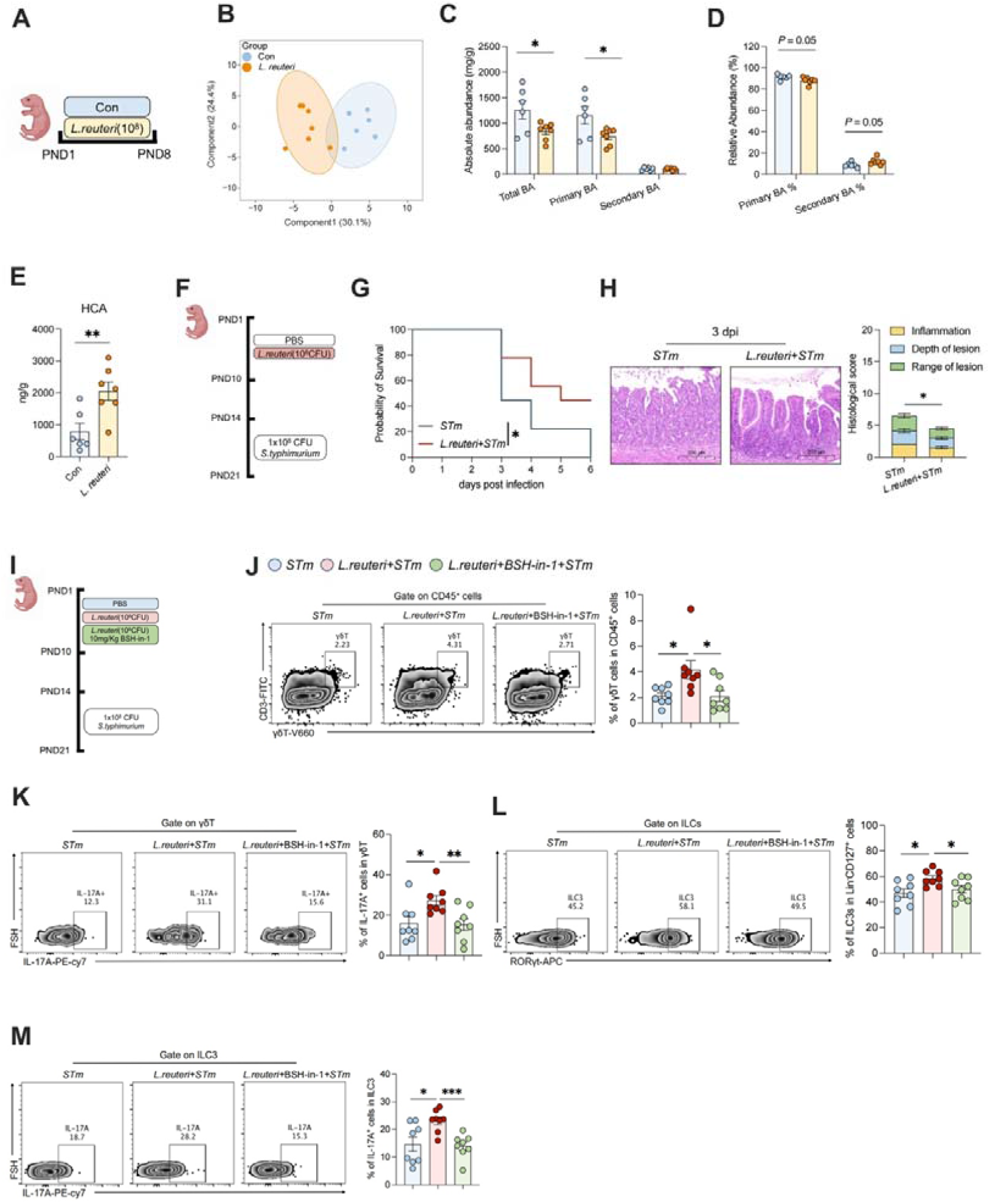
*L. reuteri* enhances type 3 immune response under *S. typhimurium* infection via the HCA-FXR-WTAP axis (A) Workflow of *L. reuteri* treatment on neonatal rats for the BAs analysis. (B OPLS-DA score plot of the ileum content BA profiles between Con and *L. reuteri* groups at 8 days. n = 6-7. (C) The absolute abundance of primary BA, secondary BA, and total BA of the ileum content BA profiles between Con and *L. reuteri* groups at 8 days. n = 6-7. (D) The relative abundance of secondary BA and primary BA of the ileum content BA profiles between Con and *L. reuteri* groups at 8 days. n = 6-7. (E) The absolute abundance of HCA between Con and *L. reuteri* groups at 8 days. n =6-7. (F) Workflow of *L. reuteri* treatment and *STm* infection on neonatal rats (G) Survival curve of the *STm* and *L. reuteri* + *STm* groups. The significance of dissimilarity was calculated by the Log-rank (Mantel-Cox) test. n = 9. (H) Representative H&E images showing histological scores in *STm* and *L. reuteri* +*STm* groups. The significance of dissimilarity was calculated by t-test. n = 6. (I) Workflow of *L. reuteri* and BSH inhibitor treatment on neonatal rats (J-K) Representative FACS plots and % of γδT and IL-17A^+^γδT in small intestine LPLs of *STm*, *L. reuteri* + *STm* and *L. reuteri* + BSH-in + *STm* groups. n = 8. (L-M) Representative FACS plots and % of ILC3 and IL-17A^+^ILC3 in small intestine LPLs of *STm*, *L. reuteri* + *STm* and *L. reuteri* + BSH-in + *STm* groups. n = 8. Data represent mean ± SEM. \**P*[<[0.05; \*\**P*[<[0.01. The difference between the two groups was analyzed by two-tailed unpaired Student’s t-test, except for (B), (G).

### *L. reuteri* enhances type 3 immune response under *S. typhimurium* infection via the HCA-FXR-WTAP axis

To investigate whether *L. reuteri* regulates FXR and m^6^A levels in the LPLs, we first examined the expression of FXR-target genes *Fgf19* and *Shp*. *L. reuteri* treatment led to a suppression of *Fgf19* and *Shp* expression in the LPLs of the rats (**Supplementary** Figure 9A). Moreover, treatment with *L. reuteri* decreased the m^6^A level and the *Wtap* level in the LPLs (**Supplementary** Figure 9B**, C**). To verify the protective function of *L. reuteri* against *S. typhimurium*, we evaluated the phenotype after *S. typhimurium* infection **(Figure 7F)**. The results showed that *L. reuteri* treatment mitigated body weight loss (**Supplementary** Figure 10A), mortality (**Figure 7G**), and diarrhea (**Supplementary** Figure 10B) after *S. typhimurium* infection. Additionally, *L. reuteri* treatment helped reduce *S. typhimurium* burden in the ileum and liver at 3 dpi (**Supplementary** Figure 10C), and ileum damage (**Figure 7H**).

We also treated with the BSH inhibitor (BSH-IN-1) based on the *L. reuteri* treatment (**Figure 7I**). As expected, the *S. typhimurium* burden in the ileum was reloaded to the level of the *STm* group after BSH-IN-1 treatment (**Supplementary** Figure 10D). Moreover, the type 3 immune response including the percentage of γδT cells and ILC3 and the ability of IL-17A secreting disappeared in the *L. reuteri* + BSH-IN-1 + *STm* group (**Figure 7J-M**). Collectively, these results demonstrated that *L. reuteri* provides protection against *S. typhimurium* infection through the type 3 immune response via the HCA-FXR-WTAP axis and depends on BSH.

## DISCUSSION

The microbiome exerts a prolonged influence on the immune system in early life such as ILCs, γδT cells, and Paneth cells [25]. In this study, we explored how the colonization of the microbiome promotes the development of intestinal type 3 immune immunity in early life. It has important clinical significance for preventing or treating bacterial infections in newborns through microbial therapy.

Although studies on germ-free models demonstrated the vital role of intestinal microbiome in type 3 immunity development [26]. Research on the relationship between microbiota and type 3 immunity is mainly limited to adult animal models. For instance, *segmented filamentous bacteria* colonize in the ileum of adult mice and promote the expansion of Th17 [27]. The previous article reported that *Akkermansia muciniphila* promotes the development of RORγt[positive innate and adaptive immune cell subsets in the early life of mice [28], whereas the underlying mechanism has not been studied clearly. Microbial metabolites broadly influence the function of type 3 immune cells in adult animals. For instance, short fatty acids facilitate the expansion of Th17 and ILC3 in the intestine via FFAR2 [9a, 29], and *Lactobacillus*- derived indole serves as a ligand for the AHR to enhance the transcription of IL-22 in ILC3 [9b]. The BAs metabolism, which is involved in the intestinal microbiome, also contributes to the function of peripheral regulatory T cells[30], ILCs [31], and intestinal stem cells [32]. Here, our research has discovered *Lactobacillus* contributes to the production of HCA and enhances the development of γδT and ILC3 through inhibition of FXR in early life, which effectively prevents intestinal bacterial infections. FXR and TGR5 are two receptors of various BAs [33], in which HCA inhibits FXR and activates TGR5 [21]. FXR deletion in ILCs promotes the IL-17 secretion of ILC3 in the intestine [22], and blocking of TGR5 reduces the IL-22 secretion of ILC3 [34]. In the current study, we found HCA enhances the transcription of *Rorc* via FXR-dependent m^6^A RNA methylation and promotes the secretion of IL- 17A in γδT and ILC3 cells to protect against the *S. typhimurium* infection. These results highlight that microbiome-derived HCA promotes intestinal type 3 immunity in early life.

The gut microbiome affects the epigenetic modification of the host [13]. Microbial-derived butyrate regulates the histone acetylation of tuft cells [35], B cells [36], T cells [37] and ILCs [38] via HDAC. In addition, GF mouse models showed that the modification of m^6^A in the intestine can be regulated by the microbiome [12]. Infection of *Fusobacterium nucleatum* in tumor cells increases the transcription of METTL3, thereby affecting the level of m^6^A modification [39]. However, research on how microbes affect the m^6^A modification of the host’s intestine is still limited. Moreover, previous studies have shown that the deficiency of the m^6^A demethylase ALKBH5 impairs ILC3 homeostasis, thereby increasing the susceptibility to *C. rodentium* infections in the intestine [40]. Our finding indicated that early-life microbiome colonization decreased the m^6^A level of intestinal LPLs via HCA. A high level of m^6^A disrupted the mRNA stability of type 3 immune factor *Rorc* in early life, and HCA enhanced the mRNA stability of *Rorc* through m^6^A modification, thereby facilitating the amplification of type 3 immune cells in the early stage, highlighting a pathway of interaction between the microbiome and immune cells.

The first step in the metabolism of secondary BAs is the hydrolysis by BSH, and *Clostridium*, *Enterococcus*, *Bifidobacterium, Lactobacillus*, and *Bacteroidetes* are all expressing BSH [41]. Our results reveal that the abundance of *Lactobacillus* is associated with secondary BA biosynthesis and the abundance of *BSH* of *Lactobacillus* is also enriched in the SB group. A recent study found that *L. reuteri* and *L. plantarum* altered BAs profiles and ameliorated dysbiosis of gut microbiome in mice [42]. In our study, we found that early-life intervention by *L. reuteri* altered BAs profiles, especially enriching the secondary BA HCA. Further, the BSH antagonist, BSH-In-1, weakened the function of *L. reuteri.* Previous studies have reported that *L. reuteri* regulates the T cells and ILC3 production of IL-17 via tryptophan and SCFAs metabolism in the colon of adult mice [43]. In the present study, we found that secondary BA biosynthesis is involved in the connection between the microbiome and type 3 immunity in the small intestine. We speculate that this is related to the physiological state, diet, and intestinal segments.

The oxygen content in the intestinal lumen is a vital factor for the colonization of the microbiome [44]. The oxygenation of colonocytes constructed by PPAR-γ signaling is beneficial to inhibiting dysbiotic *Enterobacteriaceae*, thereby ameliorating colitis [45].

Moreover, oxygen consumption also protects against pathogen bacterial colonization [46]. In early life, gut oxygen content gradually decreases with growing up [47], and the domain microbiome changes from aerobic bacteria to anaerobic bacteria [48]. Our results also show that the consumption of oxygen in the intestinal lumen can promote the microbial development of newborn rats and transform from anaerobic bacteria to facultative anaerobic bacteria dominated by *Lactobacillus*. Different from the large intestine, anaerobic bacteria such as *Bifidobacterium* cannot colonize the ileum because of the oxygen content [49]. Consequently, our data underscore the significance of rapid intestinal oxygen consumption in early-life microbial colonization.

Three types of immune responses play an important role in defending against microbial infections [50], and we found that type 3 γδT and ILC3 were elevated while Th17 did not change. We speculate that Th17 belongs to the adaptive immune cell and does not develop in neonates due to the lack of SFB [51]. While γδT and ILC3 belong to the innate immune cell and are abundant in neonatal mice [4, 6c]. IL-17A is produced early after infection by pathogens, which indicates the presence of innate pathways of IL-17 secretion. For example, the expression of IL-17 was found to occur 5 hours after the inoculation of rhesus macaque ileal loops and 48 hours after oral infection of mice with *Salmonella* [52]. We also found that the *Il17a* was elevated 24h after *S. typhimurium* infection, and *S. boulardii* promoted the secretion of IL-17A in the small intestine but not IL-22. Further, the high level of *Il17a* in the intestine presents lower *S. typhimurium* loading. Collectively, IL-17A-producing type 3 γδT and ILC3 play a crucial role in protecting against *S. typhimurium* infection in early life.

This study has some limitations regarding mechanism investigation and clinical translation. Though we found that *Lactobacillus* contributes to the production of HCA depending on the BSH, the total metabolite pathway of HCA is still unknown. Other bacteria may interact with *Lactobacillus* to produce HCA together. Further research like bacterial co-culture, and multi-omics research is needed to investigate the underlying mechanisms. Additionally, in addition to *RORC*, there may be other genes involved in the regulation of ILC3 and γδT cells as target genes of WTPA. Studies conducted at the genome-wide level by meRIP-seq may provide additional valuable evidence, and the application of genetically engineered animals is also helpful in the verification of mechanisms.

Taken together, our research shows that a hypoxic environment in the intestine promotes the colonization of the microbiome during early life, particularly *Lactobacillus* species. The HCA derived from *Lactobacillus* suppresses FXR signaling and enhances the stability of *Rorc* through the FXR-WTAP-m^6^A axis, which leads to the expansion of type 3 γδT and ILC3 and increased resistance to *S. typhimurium* infection in the intestine.

## Materials and Methods

### Animal experiments

Animal welfare and experimental procedures were monitored according to the Guide FOR THE CARE AND USE OF LABORATORY ANIMALS (Eight Edition) and were approved by the ethical committee at Huazhong Agricultural University, ID Number: 202401050001. Wistar rats were obtained from the Animal Experiment Center at Hubei Disease Prevention and Control Center (Wuhan, China). The Rats were housed under specific pathogen-free conditions in a temperature-controlled room at 23 ± 2[. There was free access to food and water.

For the *S. boulardii* intervention, pregnant Wistar rats were chosen and monitored daily until parturition. After birth, each litter was culled to a size of 12 pups. On postnatal day 1 (PD1), pups were randomly divided into two groups (PBS or *S. boulardii*) in each litter. Rats in the PBS groups were orally administrated with PBS once per day from PD1 to PD10; Rats in the SB groups were orally administrated with 1 × 10^9^ colony-forming units (CFU) of *S. boulardii* once per day from PD1 to PD10. The test product was Levucell SB® (Lallemand SAS, Blagnac, France). In PD3, PD5, PD8, PD11, and PD14, 2 rats from every litter from two groups were randomly chosen and were asphyxiated by CO_2_ inhalation and sacrificed.

For early-life *S. typhimurium* infection, in PD14, rats were infected with 1 × 10^8^ CFU/mL of *S. typhimurium* (strain SL1344), which was provided by Hangzhou Yinyuan Biotechnology Company. The body weight and survival were recorded daily. Meanwhile, the diarrhea status was recorded daily according to the diarrhea score.

The protocol of WIMT was appropriately modified based on Li et al [20] and Kim et al [46b]. Briefly, choose 8-day years old rats, and extract the content of jejunum, ileum, cecum, and colon in the anaerobic chamber (LABIOPHY, Dalian, China) with 10% CO_2_, 10% H_2_, and 80% N_2_. 40 mL PBS with 10% glycerol added to 1g of content and vortexed, then passed through a 70 μm filter to remove large particulate, next centrifuged at 200 × g for 2 min, and immediately dispensed the supernatant to cryotubes, and stored at –80°C.

HCA (Merck, Cat#: 700159P) intervened at 100 mg/Kg/d, GW4064 (MCE, Cat#: HY- 50108) and BSH-IN-1 (MCE, HY-135659) intervened at 10 mg/Kg/d. *L. reuteri* strain (BNCC 186563) was cultured in Murashige and Skoog (MRS) medium (Hopebiol, Cat#: HB0384-1) for 24 h at 37 °C. Collected the *L. reuteri* at the logarithmic phase (6 h), diluted to different concentrations for use.

Rats were asphyxiated by CO_2_ inhalation and sacrificed, intestinal tissues were collected from all rats and fixed in 4% paraformaldehyde fix solution, and intestinal tissues were collected and stored at −80 [ for other tests.

### Hypoxyprobe

Hypoxyprobe (Cat#: HP1) was purchased from Hypoxyprobe, Inc (Burlington, UAS). The method and does of use according to the Instruction manual. Briefly, 60mg/Kg of Hypoxyprobe was intraperitoneally injected in rats, and the tissues were harvested after 1.5 h and then immunofluorescence staining was done.

### RNA isolation and quantitative real-time PCR

For LPLs and tissue, total RNA was extracted using the TRIzol reagent (Vazyme, Cat#: R401-01). For sorted cells, total RNA was extracted using the PureLink™ RNA Micro Scale Kit (Thermo, Cat#: 12183016). Then detected the concentration of RNA by NanoDropfi ND-1000 Spectrophotometer (Thermo, Waltham, USA) and transcribed into cDNA using Reverse Transcriptase (ABclonal, Cat#: RK20433). Quantitation of the mRNA levels by quantitative real-time (qPCR) was performed on a real-time PCR system (Bio-Rad, California, USA) using SYBR Green qPCR Master Mix (Vazyme, Cat#: Q321). The sequences of the primers are listed in **Supplementary Table 1**

### Intestinal microbiome sequence

The method previously described by Xia et al [53]. Briefly, a QIAamp DNA Stool Mini Kit (QiagenLtd, Frankfurt, Germany) was used to extract the total microbial genomic DNA from intestinal contents following the manufacturer’s instructions. Amplicon libraries were sequenced on an Illumina MiSeq (Illumina, Santiago, USA) at the Personalbio, Shanghai, China. QIIME 2 2019.4 was used to perform the microbiome bioinformatics [54]. DADA2 plugin was used to filter quality, denoise, merge and remove chimera for sequences [55]. To assign taxonomy to ASVs, the classify-sklearn naïve Bayes taxonomy classifier in the feature-classifier plugin was used [56] against the Greengenes 13_8 99% operational taxonomic unit reference sequences [57]. The Chao1, Shannon, Simpson, and Observed species values were used to assess the diversity of Alpha. Beta diversity measures depended on Bray- Curtis distance.

### Isolation of intestinal LPLs and flow cytometry

The isolation of intestinal LPLs was performed as previously described with probable modification [58]. Briefly, the small intestine was dissected, and fat and mesentery tissues were removed. Intestines were dissected longitudinally and subsequently cut into several pieces, followed by washing with cold Hanks’ Balanced Salt Solution (Solarbio, Cat#: H1046). To remove epithelial cells, the intestines were then incubated successively with 2% FBS, 1 mM dithiothreitol (Servicebio, Cat#: GC205010), 30 mM EDTA (Sanangon, Cat#: B540625)-PBS once for 30 min at 37 °C while shaking at 200 rpm. The tissues were then digested with 50 μg/mL DNase I (Roche, Cat#: 10104159001) and 300 U/mL collagenase VIII (Sigma, Cat#: C2139) in RPMI1640 medium (Servicebio, Cat#: G4533) for 30 min at 37 °C while shaking at 200 rpm. The digested tissues were homogenized by vigorous shaking and filtered with a 70 µm cell strainer. Mononuclear cells were then harvested from the interphase of an 80 and 40% Percoll (Cytiva, Cat#: 365237) gradient after spinning at 800×g for 15 min at RT.

For cytokine staining, cells were stimulated with Cell Stimulation Cocktail (plus protein transport inhibitors) (500X) (eBioscience, Cat#: 00-4975-93) for 5 h. The panel design refers to ref [59], the detailed antibodies used are shown in **Supplementary Table 2**. Analysis was performed on a CytoflexLX (Beckman, USA), and sorting was performed on a CytoFLEX SRT (Beckman, USA). FACS-gating strategies for different immune cell populations are shown in **Supplementary Figure1**

### Hematoxylin and eosin (H&E) staining and Immunofluorescence staining

For H&E staining, deparaffinize, and hydrate to water. Secondly, stain in hematoxylin solution: immerse slides in hematoxylin solution for 3 to 5 min and rinse them in water. Then differentiate sections with acid alcohol, and rinse again. Blue up sections with ammonia solution, and wash in slowly running tap water. Thirdly, stain in eosin. Last, dehydrate and mount. The section was scanned with Pannoramic SCAN (3DHISTECH CaseViewe, Budapest, Hungary). The morphology and histological Score were analyzed by CaseViewer software (3DHISTECH CaseViewe, Budapest, Hungary). We assessed the distance from the apical side to the basal side of the crypt based on at least 10 intact and well-oriented crypts of each sample. We assessed the histological score based on the degree of inflammation (1-2), lesion depth (1-4), and lesion extent (1-4).

For Immunofluorescence staining, tissue sections were washed 3 times with PBS. Block with 5% BSA for 1 h at room temperature. Add the PMDZ primary antibody diluent (1:100 in PBS) according to the purpose of the experiment and place it in a refrigerator at 4°C overnight. The primary antibody was recovered the next day and washed three times by adding PBS. Then wash with PBS three times in the dark. Then add DAPI staining solution 200 uL at room temperature for 20 min in the dark. Remove the above staining solution and add PBS to wash three times in the dark. Prepare the slides, add 5 uL of an anti-quenching mounting solution to each sample, put the slides upside down on the mounting solution, and store them at 4°C in the dark. After overnight observation under a laser confocal microscope, be photographed to record the results.

### Bacterial Load Assay

To quantitatively determine the bacterial load. 50 mg of the sample was resuspended in a sterile EP tube with 0.5 mL of PBS with 1% Triton X-100 (Servicebio, Cat#: GC204003) and recorded 10^−1^. We added 20 μL of the suspension to 200 μL of PBS and recorded 10^−2^. This was followed by plating onto Bismuth Sulfite Agar (Huankai, Cat#: 027319) using serial dilutions. After incubation for 18 h at 37[, the counting of CFU occurred.

### Western blot

The detailed protocol was described in the ref [60]. Briefly, tissues were extracted with RIPA Lysis Buffer supplemented with 100× Protease Inhibitor Cocktail. The protein concentration was determined using the BCA Kit. Proteins were separated on an SDS gel followed by blotting on PVDF membranes. The membrane was blocked for 1 h, and then incubated with the primary antibody overnight at 4[: anti-β-actin (1:50000, ABclonal, Cat#: AC026), anti-Claudin-1 (1:1000, Affinity, Cat#: AF0127), anti-Occludin (1:1000, Affinity, Cat#: DF7504). Secondary antibodies, anti-mouse, or rabbit IgG-HRP (1:5000, ABclonal, Cat#: AS014) were used to detect primary antibodies. Binding was detected using an enhanced chemiluminescence detection kit, and densitometry was performed using Image-J software (Bethesda, Maryland, USA).

### Elisa detection

Tissues were pulverized with a homogenizer. Then the supernatant was collected for the Elisa assay. Before the Elisa assay, the protein concentration was determined using the BCA Kit. Rat ELISA kits including IL-22 (MM-0670R1) and IL-17A (MM- 70049R1) were purchased from Jiangsu Meimian Industrial Co., Ltd to determine the concentrations of cytokines in the intestine according to the manufacturer’s instructions.

### Molecular docking

The crystal structures of the complex of the FXR ligand binding domain were downloaded from the RCSB Protein Data Bank (PDB ID: 5YXJ). The molecular structure of HCA was retrieved from PubChem Compound (CID: 92805). prepared by SYBYL-X 2.0. Molecular docking studies were performed by AutodockVina 1.2.2, a silico protein–ligand docking software was employed. The binding interaction was edited using PyMOL.

### Cell culture

HEK293T cell cultured high glucose Dulbecco’s modified Eagle’s medium (DMEM) (Hyclone, Cat#: SH30243.01), containing 10% fetal bovine serum (NEWZERUM, Cat#: FBS-S500) and 1% penicillin/streptomycin (Hyclone, Cat#: SV30010) at 37°C under a 5% CO_2_ atmosphere. HEK293 cells were transfected with different vectors and exposed to different concentrations of HCA (Merck, Cat#: 700159P) and GW4064 (MCE, Cat#: HY-50108) for 24 h used for luciferase reporter assay.

For HCA treatment, freshly isolated intestinal LPLs were cultured in the RPMI 1640 medium (servicebio, Cat#: G4533) containing 10% fetal bovine serum (NEWZERUM, Cat No: FBS-S500) and 1% penicillin/streptomycin (Hyclone, Cat#: SV30010), and Cell Stimulation Cocktail (plus protein transport inhibitors) (500X) (eBioscience, Cat#: 00-4975-93) at 37°C under a 5% CO_2_ atmosphere, with different concentrations of HCA as indicated in the text in a 24-well plate for 16 h and then collected for the following analysis.

### Luciferase reporter assay

pGL3-Basic-WTAP firefly luciferase reporter vector and different fragments of the WTAP promoter were constructed by Tsingke Biotechnology. The human NR4H1 (FXR) expression vector and human ASBT expression vector were purchased from GeneCopoeia. HEK293T cells were cultured and co-transfected with the human FXR expression vector, human ASBT expression vector, pGL3-Basic-WTAP firefly or WTAP P1-P4, deletion, and mutation luciferase reporter vector, and the Renilla luciferase control vector (Promega, Madison, WI) using Lipo8000™ transfection reagent (Beyotime, Cat#: C0533). Luciferase assays were performed by Dual-Glo^®^ Luciferase Assay System (Promega, Cat#: E2920). Firefly and Renilla luciferase activities were measured by Synergy2 (BioTek, USA).

### BAs analysis

BAs contents were detected by MetWare ( http://www.metware.cn/) based on the AB Sciex QTRAP 6500 LC-MS/MS platform. Briefly, Samples (20 mg) were extracted with 200 μL methanol/acetonitrile(v/v=2:8) after the samples were grinded with a ball mill. 10 μL internal standard mixed solution (1 μg/mL) was added into the extract as internal standards (IS) for the quantication. Put the samples at -20°C for 10 min to precipitate protein. Then centrifugation for 10 min (12000 r/min, and 4°C), the supernatant was transferred to clean plastic microtubes. The extracts were evaporated to dryness and reconstituted in 100 μL 50% methanol (V/V) for further analysis. The sample extracts were analyzed using an LC-ESI-MS/MS system.

### m^6^A Dot Blot

The dot blot refers to the previous study. First, prepare a denaturing solution, which consists of 60 μL 20xSSC buffer (Biosharp, Cat#: BL164A) and 40 μL 37% deionized formaldehyde to create 100 μL RNA denaturing solution. Mix the RNA and denaturing solution at a 1:1 ratio and then denature the nucleic acids in the solution by heating them in a PCR instrument at 95°C for 5 min. Next, quickly spot the denatured samples onto a nitrocellulose (NC) membrane, and then cross-link them under a 302 nm UV lamp for 6 minutes. Detect the cross-linked NC membrane using a specific m^6^A antibody (Abclonal, Cat#: A19841) as described in the Western Blot method.

### RNA stable ability assay

The sorted cells were cultured in RPMI 1640 medium (Servicebio, Cat#: G4533) supplemented with 10% fetal bovine serum (NEWZERUM, Cat No: FBS-S500) and 1% penicillin/streptomycin (Hyclone, Cat#: SV30010) at 37°C under a 5% CO_2_ atmosphere. Actinomycin D (MCE, Cat No: HY-17559) was added to a final concentration of 1 μM, and cells were harvested at t[=5 h after actinomycin D treatment. The RNA was extracted and subjected to RT-qPCR analysis.

### The SELECT detection assay

Epi-SELECT m^6^A site identification kit (Epibiotek, Cat#: R202106M-01) was purchased from Guangzhou Epibiotek Co., Ltd.. The detail method was referred to by Xiao et al [61]. Briefly, 20 ng RNA was mixed with Up Primer, Down Primer, and dNTP in 1× CutSmart buffer, next annealed and extended in the Thermal Cycler (Bio- Rad, California, USA). Subsequently, the mixture containing SELECT™ DNA polymerase, SELECT™ ligase, and ATP was added to the former mixture, next conducted single base extent in the Thermal Cycler (Bio-Rad, California, USA). Last, qPCR was performed on a real-time PCR system (Bio-Rad, California, USA) using SYBR Green qPCR Master Mix (Vazyme, Cat#: Q321). Primer sequences for specific m^6^A sites of *Rorc*: Rorc_up_probe: tagccagtaccgtagtgcgtgCTTCTGGGTGCTTGCCACCAG, Rorc_down_probe: 5phos/ CTCTGAGCTAGATCCATCTCCCCCcagaggctgagtcgctgcat.

### Statistical analysis

Data were analyzed using GraphPad Prism (v 9.0.0) and Microsoft Excel (2023), flowjo (v 10.4), and expressed as the means ± standard error of the mean (SEM). Firstly, data were assessed on the normality of the distribution by the Shapiro−Wilk test. Then, a t-test was used to analyze the data with a normal distribution, and the data with a non-normal distribution were analyzed by the Mann−Whitney U-test. Statistical significance was defined as *P* < 0.05. The Random Forest algorithm was used to model the maturation process of intestinal microbiome, modified as appropriate refer as Gao et al [62]. An unsupervised clustering algorithm was used to recognize the mature and immature microbiome, modified as appropriate refer as Zhou et al [63]. For the Co-occurrence network diagram, the differences in the abundance were analyzed by Spearman’s rank correlation analysis, the ASVs were selected with presence in >30%. Co-occurrence networks were constructed using data with correlation coefficients |r-value| > 0.8 and P-values < 0.05. Gephi (v 0.10.1) was used for visualization and network analysis. Graphical representations were used by GraphPad Prism version 9.0.0 software (San Diego, California, USA), imageGP [64]. Correlation analysis was performed using Spearman’s rank test correlation analysis. The Graphical Abstract was conducted by Figdraw.

## Acknowledgments

This research was supported by the Hubei Province Science and Technology Innovation Major Project (2022BBA0012), Fundamental Research Funds for the Central Universities (2662022DKPY005, 2662022DKPY002), China Agriculture Research System of MOF and MARA, Technical Team for Genetic Improvement and Healthy Breeding of Livestock and Poultry (2021-620-000-001-030).

## Author contributions

Z.P.Y., P.J., and H.K.W. conceived the project and designed the study. Z.P.Y., Z.Y.L, Y.J.Y, M.Z., N.G. X.C.L, X.R.L performed the experiments and analyzed the data. Z.P.Y., P.J., and H.K.W. wrote and revised the manuscript. All the authors edited the manuscript and approved the final manuscript.

## Declaration of interests

The authors declare no competing interests

---

**Supplementary Figure 1.**
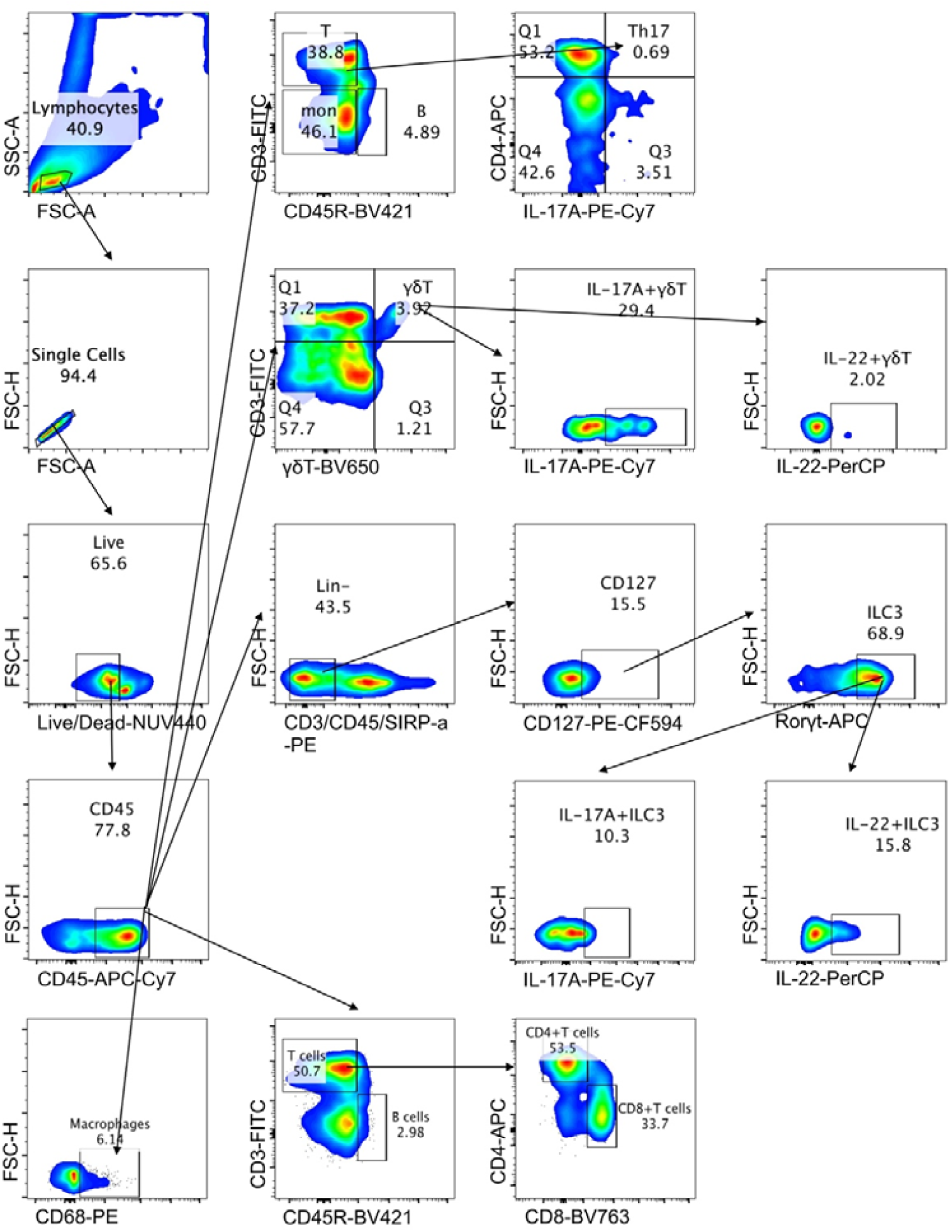
FACS gating strategies (A) Representative FACS gating strategies used for the identification of different immune cell populations in this study

**Supplementary Figure 2.**
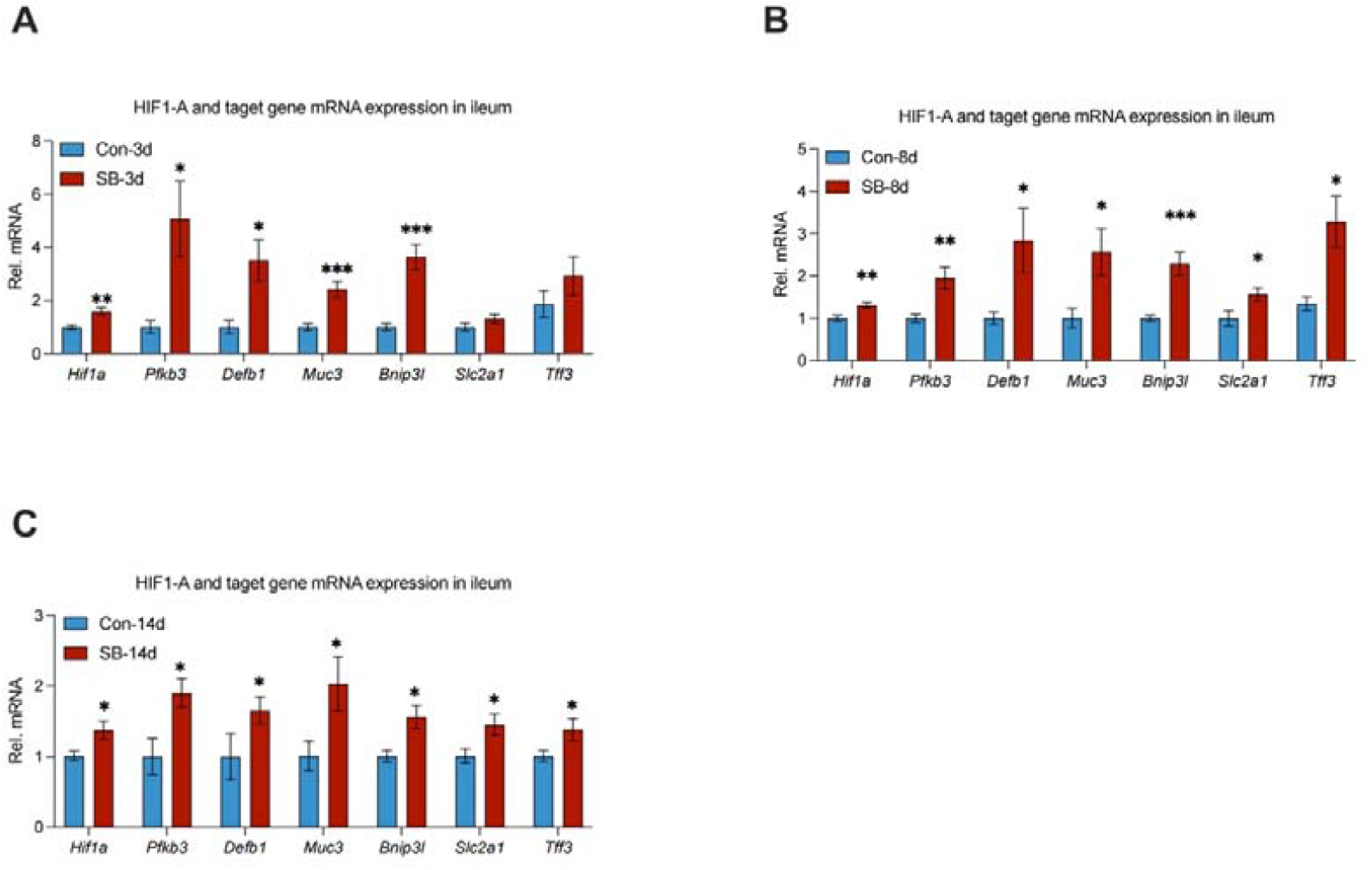
Early intervention of *S. boulardii* contributes to intestinal hypoxia in early life (A-C) Relative mRNA levels of *Hif1a* and target genes in ileum tissues of rats, including *Pfkb3*, *Defb1*, *Muc3*, *Bnip3l*, *Slc2a1*, and *Tff3*. n = 8-12. Data represent mean ± SEM. \**P*[<[0.05; \*\**P*[<[0.01; \*\*\**P*[<[0.001. The difference between rats after PBS and *S. boulardii* treatment was analyzed by two-tailed unpaired Student’s t-test.

**Supplementary Figure 3.**
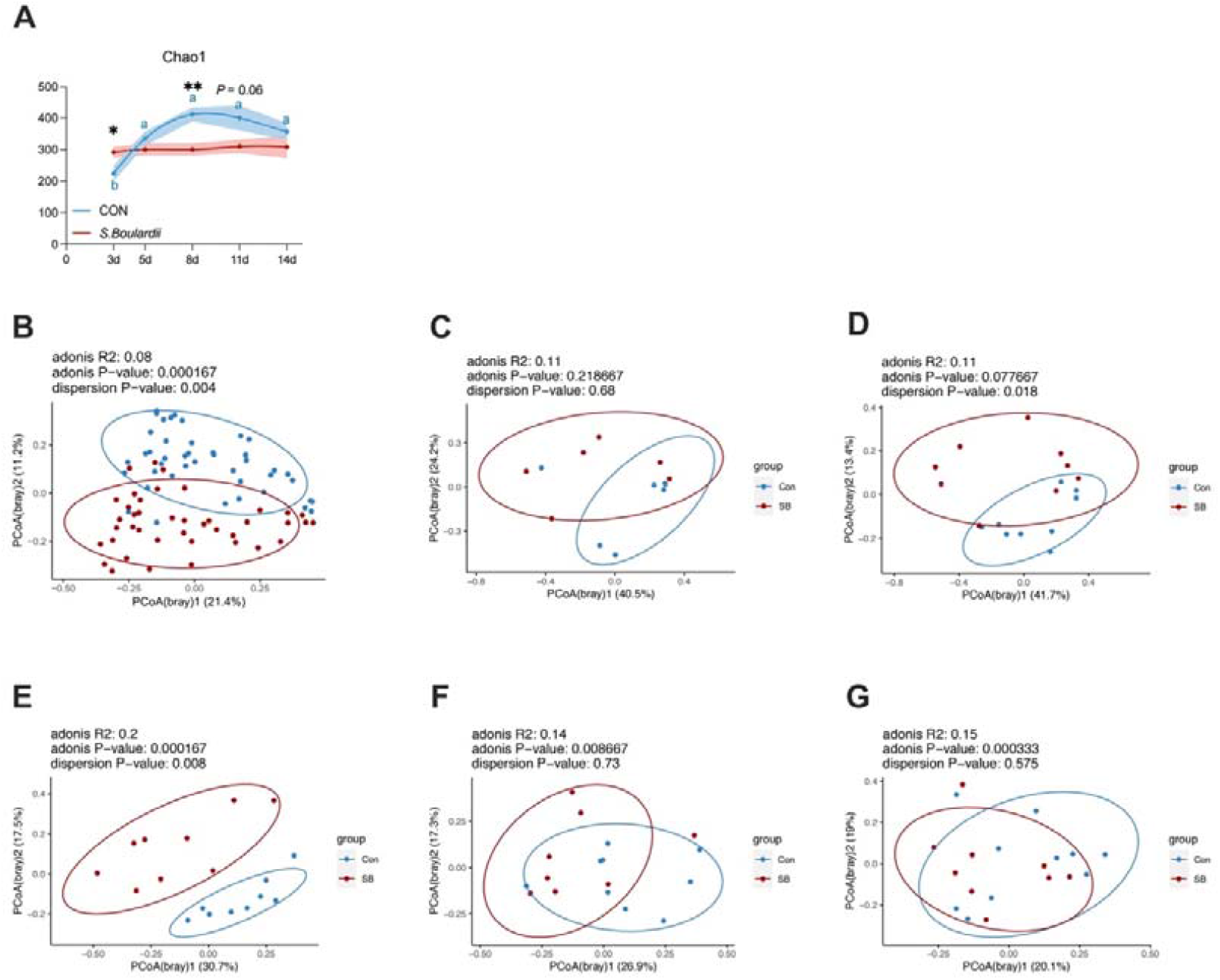
*S. boulardii*-induced oxygen consumption changes the composition of the microbiome (A) The Chao1 index of ileum content microbiome. Different blue letters represent significant differences (*p* < 0.05) between different ages in the Con group based on a one-way ANOVA test. The difference between the Con and SB groups was analyzed by two-tailed unpaired Student’s t test. (B) PCoA (principal coordinate analysis) plot of rats from the Con and SB group on the Bray-Curtis. The significance of dissimilarity was calculated by adonis and dispersion. (C-G) PCoA (principal coordinate analysis) plot of rats from the Con and SB group on the Bray-Curtis at 3, 5, 8, 11, 14 days. The significance of dissimilarity was calculated by adonis and dispersion.

**Supplementary Figure 4.**
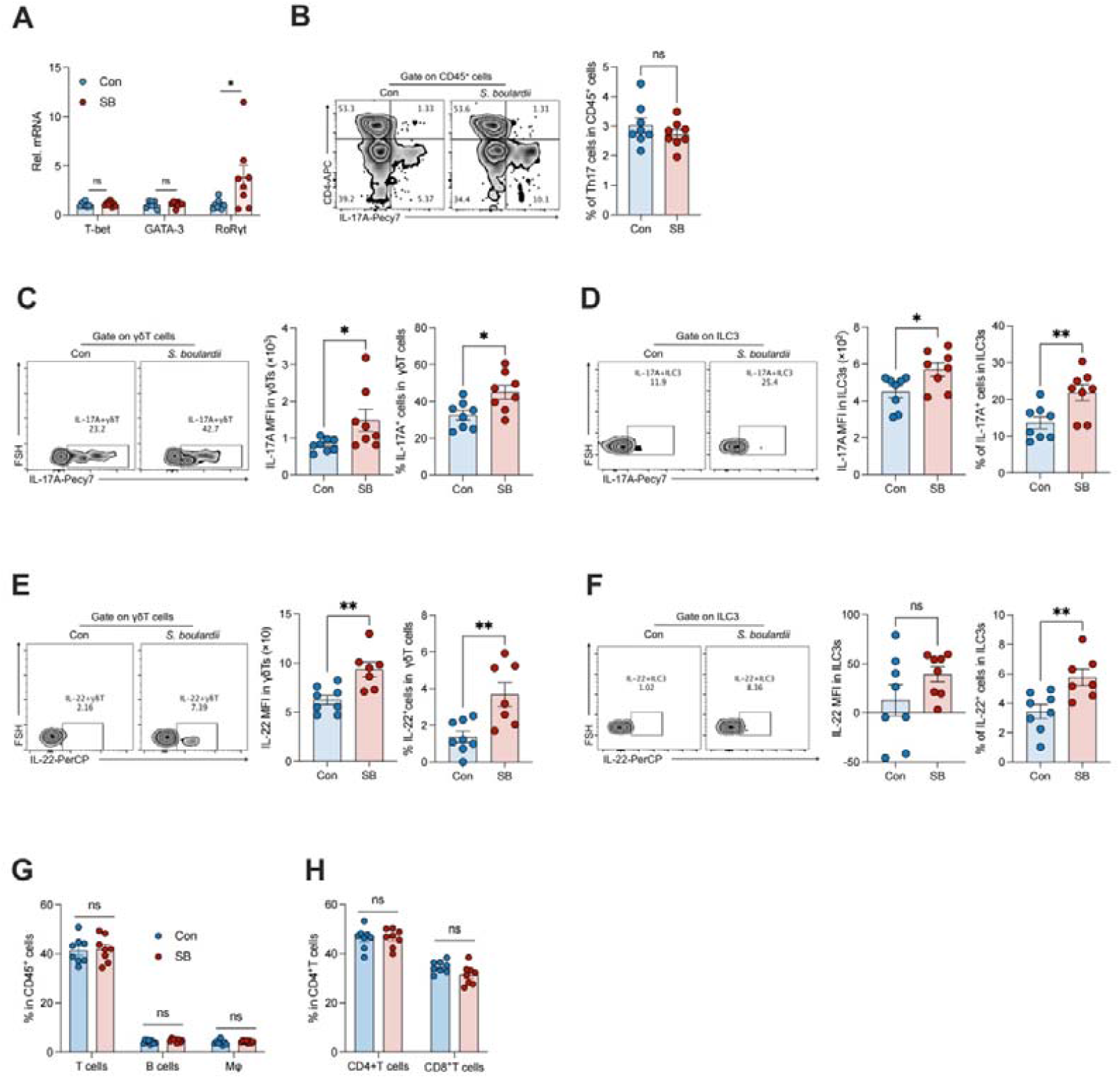
*S. boulardii*-induced oxygen consumption facilitated the maturation of type 3 immune cells (A) Relative mRNA levels of T-bet, GATA-3, and RoRγt in small intestine LPLs of rats. n = 8. (B) Representative FACS plots and % of Th17 in small intestine LPLs of rats. n = 8. (C-D) Representative FACS plots and % of Th17, γδT, and ILC3 in small intestine LPLs of rats. n = 8. (E-F) Representative FACS plots and % of IL-22^+^ γδT and ILC3 in small intestine LPLs of rats. n = 8. (G) FACS analysis of T cells, B cells, Macrophages. n = 8. (H) FACS analysis of CD4^+^T cells, and CD8^+^T cells. n = 8. Data represent mean ± SEM. \**P*[<[0.05; \*\**P*[<[0.01; ns, no significance. The difference between two groups was analyzed by two-tailed unpaired Student’s t-test.

**Supplementary Figure 5.**
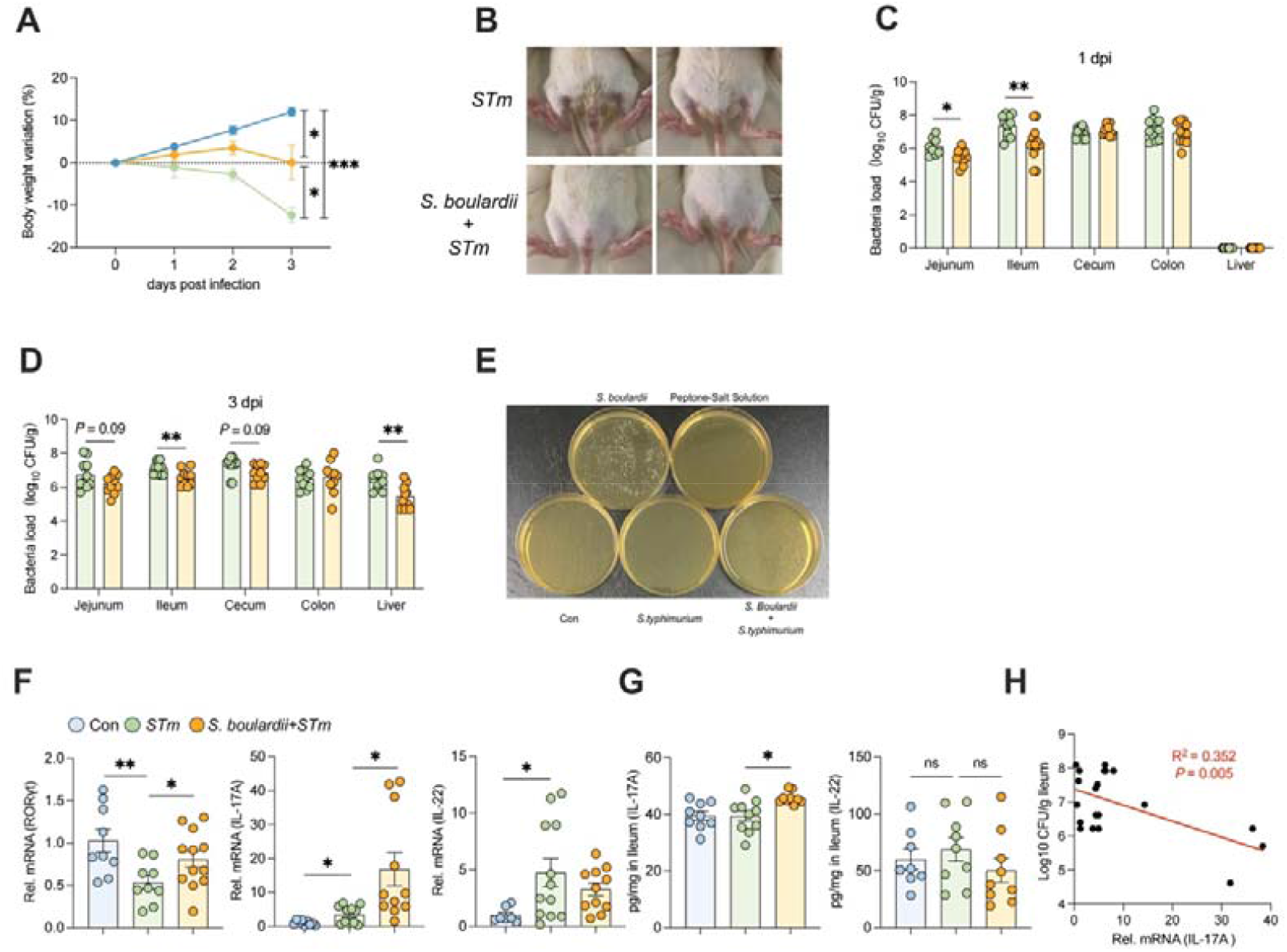
*S. boulardii*-induced oxygen consumption confers protection from *S. typhimurium* infection (A-B) Bodyweight variation and representative diarrhea images of Con, SB, and SB + *STm* groups. n = 6. (C-D) *STm* burden in the Jejunum, Ileum, Cecum, Colon, Liver at 1 dpi (F) and *STm* burden in the Jejunum, Ileum, Cecum, Colon, Liver at 3 dpi (G). n = 10 - 12. (E) *S. boulardii* burden in the ileum content at 1 dpi. (F) RT-qPCR results of mRNAs encoded by the type 3 immune factors *Rorc*, *Il17a*, *Il22* in ileum tissues of rats in Con, SB, and SB + *STm* groups. mRNA levels were normalized as to β-actin. n = 10-12. (G) Elisa results of protein of IL-17A and IL-22 in ileum tissues of rats in Con, SB, and SB + *STm* groups. n = 10-12. (H) Correlation between *STm* burden with mRNA expression of *Il17a* in ileum tissues of rats. Data represent mean ± SEM. \**P*[<[0.05; \*\**P*[<[0.01; ns, no significance. The difference between two groups was analyzed by two-tailed unpaired Student’s t-test, except for (H).

**Supplementary Figure 6.**
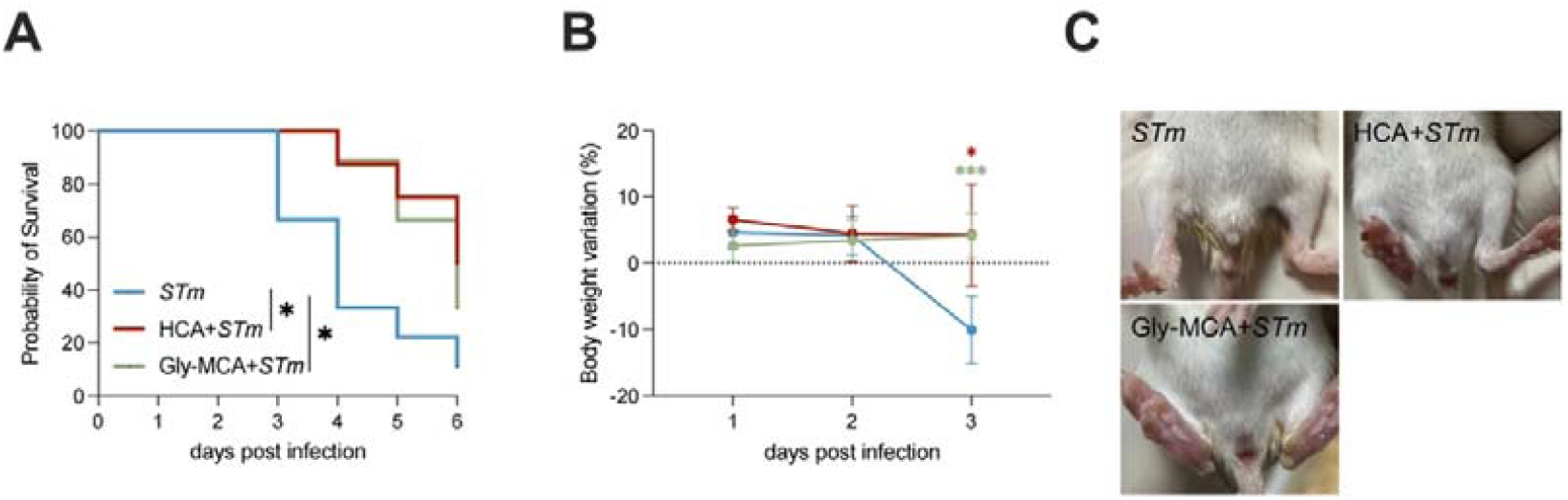
HCA and Gly-MCA enhance the protection from *S. typhimurium* infection (A) Survival curve of the *STm*, HCA + *STm*, Gly-MCA + *STm* groups. The significance of dissimilarity was calculated by the Log-rank (Mantel-Cox) test. n = 8- 9. (B) Bodyweight variation of the *STm*, HCA + *STm*, Gly-MCA + *STm* groups. The significance of dissimilarity was calculated by two-tailed unpaired Student’s t-test. n = 8-9. (C) Representative diarrhea images of the *STm* and *L. reuteri* + *STm* groups.

**Supplementary Figure 7.**
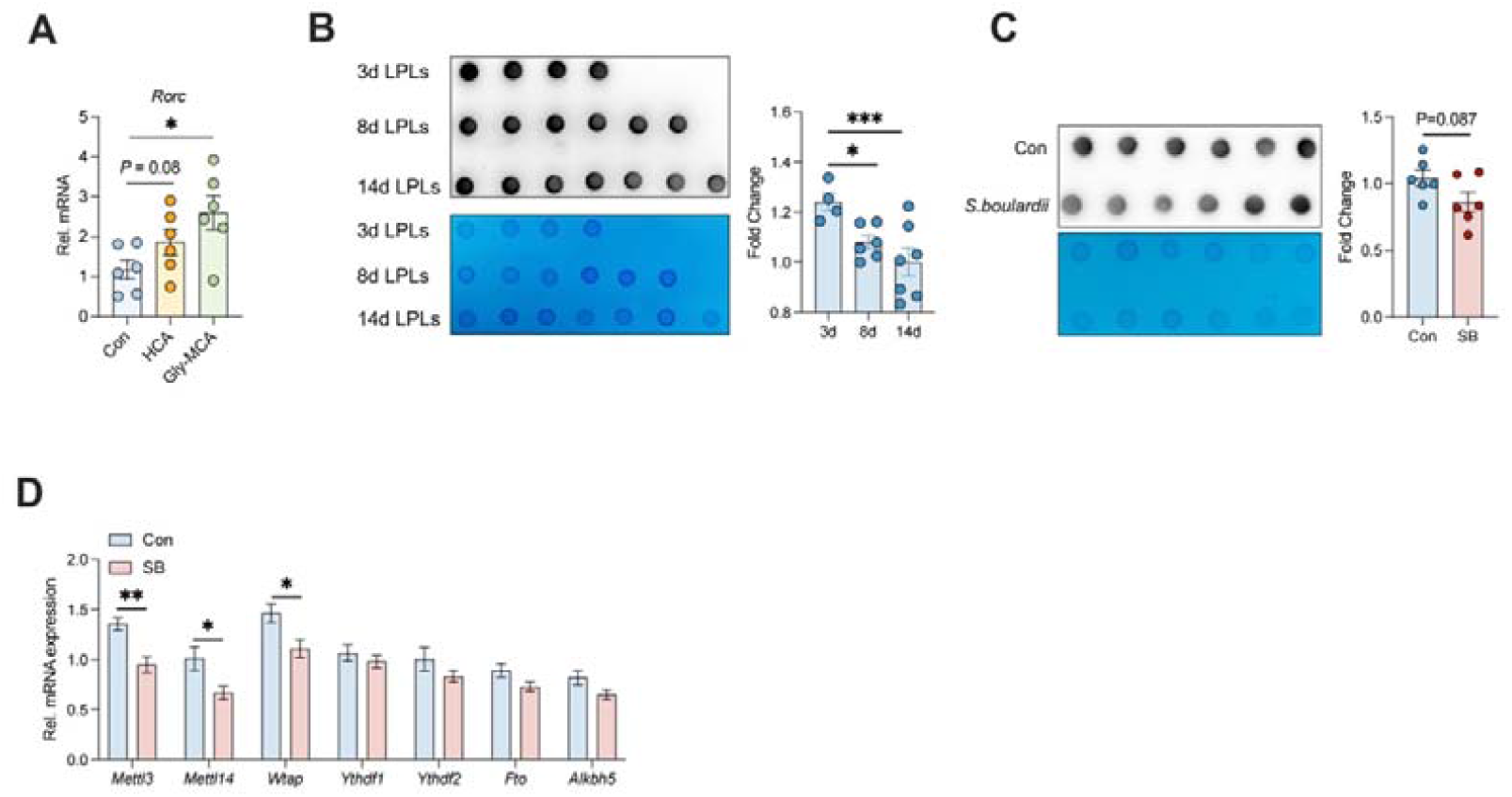
HCA affects the transcription of *Rorc* via m^6^A (A) RT-qPCR results of mRNAs encoded by *Rorc* in LPLs isolated from rats from Con, HCA, and Gly-MCA groups, n = 6. (B) Dot blot to detect the m^6^A level of mRNA isolated from total RNA of LPLs isolated from 3, 8, and 14-day-old rats, n = 4-7. (C) Dot blot to detect the m^6^A level of mRNA isolated from total RNA of LPLs isolated from rats from Con and SB groups, n = 6. (D) RT-qPCR results of mRNAs encoded by the m^6^A relative gene *Wtap, Mettl3, Mettl14, Ythdf1, Ythdf2, Fto, and Alkbh5* in LPLs isolated from rats from Con and SB groups, n = 8. Data represent mean ± SEM. \**P*[<[0.05; \*\**P*[<[0.01; \*\*\**P*[<[0.001; ns. The difference between two groups was analyzed by two-tailed unpaired Student’s t-test.

**Supplementary Figure 8.**
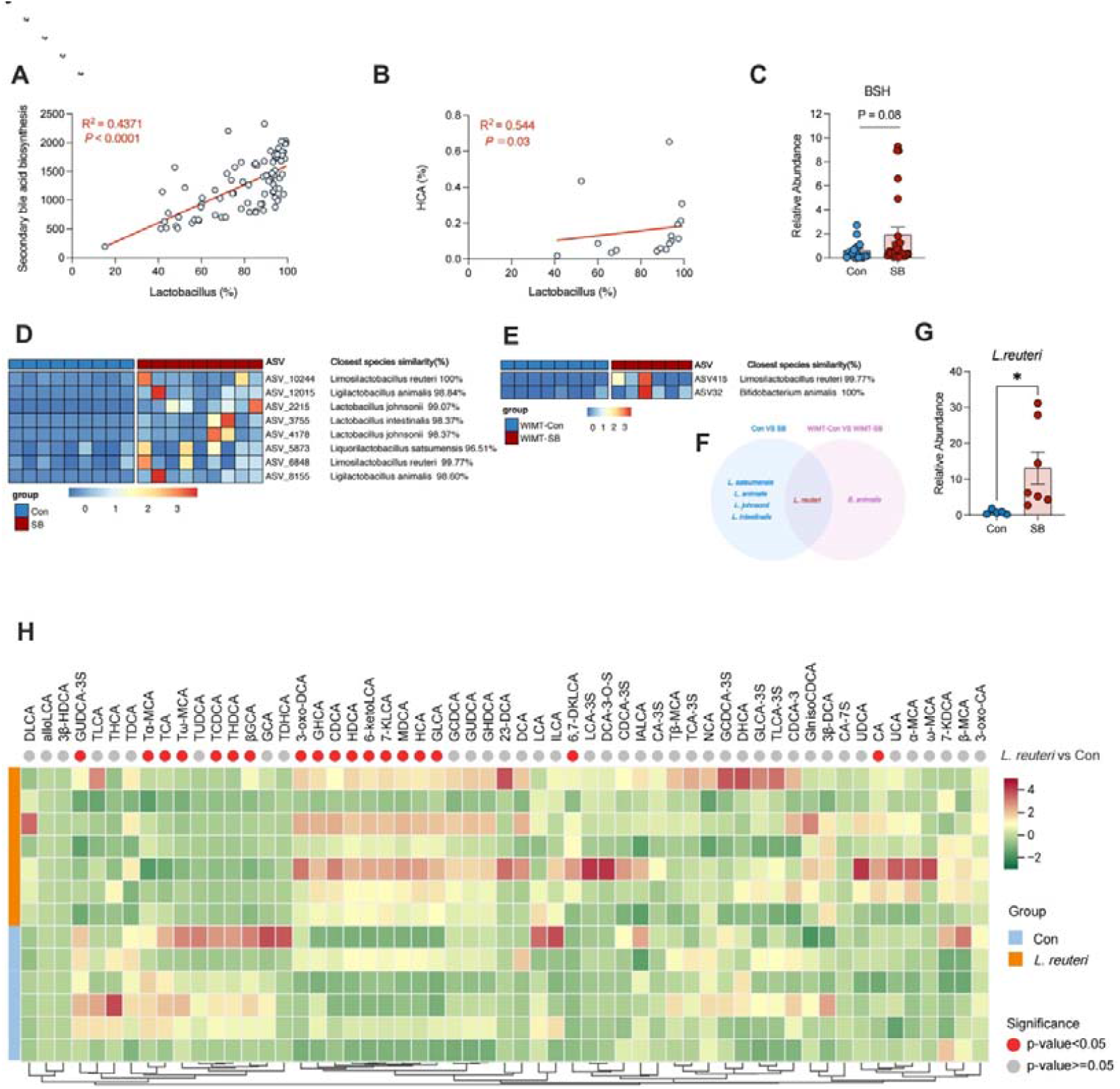
Oxygen consumption enriched *L. reuteri* changed BAs profiles (A) Correlation between the relative abundance of *Lactobacillus* with secondary BA biosynthesis in ileum content of rats. (B) Correlation between the relative abundance of *Lactobacillus* with the relative abundance of HCA in ileum content of rats. (C) RT-qPCR results of relative abundance of BA hydrolase (*bsh*) in ileum content of Con and SB rats (D and E) Heatmap showing the significantly different relative expression levels of ASVs in Con-SB rats and WIMT-PBS-SB rats. The ASVs sequences were compared at the EzBioCloud. (G) RT-qPCR results of relative abundance of *L. reuteri* in ileum content of Con and SB rats. n = 5-7. (H) Heatmap showing the absolute levels of BAs between Con and *L. reuteri* groups. Red dot presents *p*-value < 0.05, gray dot presents *p*-value >: 0.05. n = 6-7. Data represent mean ± SEM. \**P*[<[0.05. The difference between the two groups was analyzed by two-tailed unpaired Student’s t-test, except for (A), (B).

**Supplementary Fig 9.**
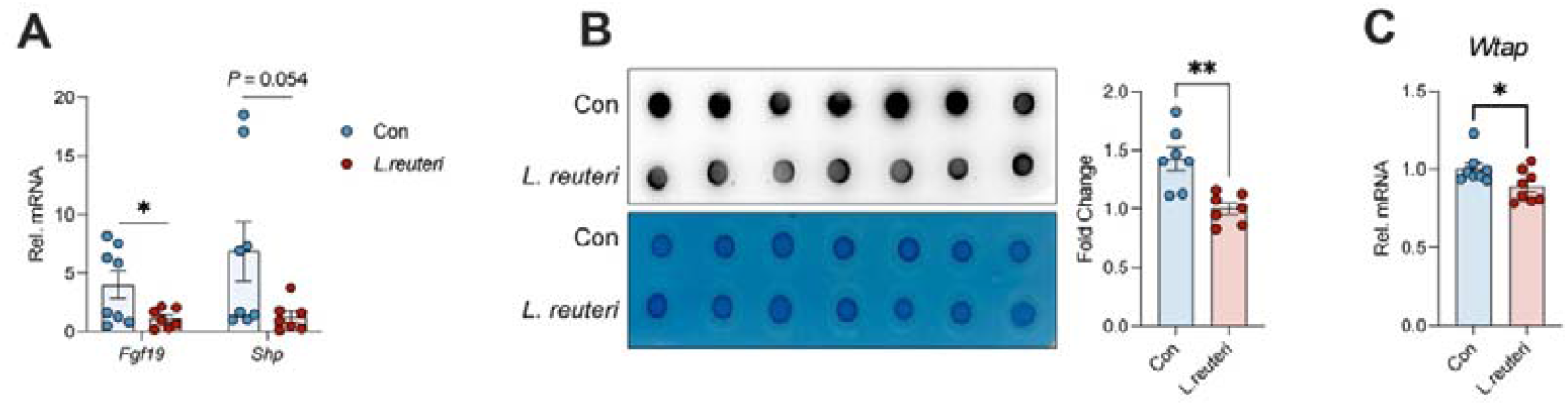
*L. reuteri* affects the FXR-WTAP-m^6^A axis (A) RT-qPCR results of mRNAs encoded by the FXR target gene *Fgf19* and *Shp* in LPLs of rats between Con and *L. reuteri* groups, n = 8. (B) Dot blot to detect the m^6^A level of mRNA isolated from total RNA of LPLs isolated from rats from Con and *L. reuteri* groups, n = 7. (C) RT-qPCR results of mRNAs encoded by *Wtap* in LPLs of rats between Con and *L. reuteri* groups, n = 8. Data represent mean ± SEM. \**P*[<[0.05; \*\**P*<0.01. The difference between the two groups was analyzed by two-tailed unpaired Student’s t-test.

**Supplementary Fig10.**
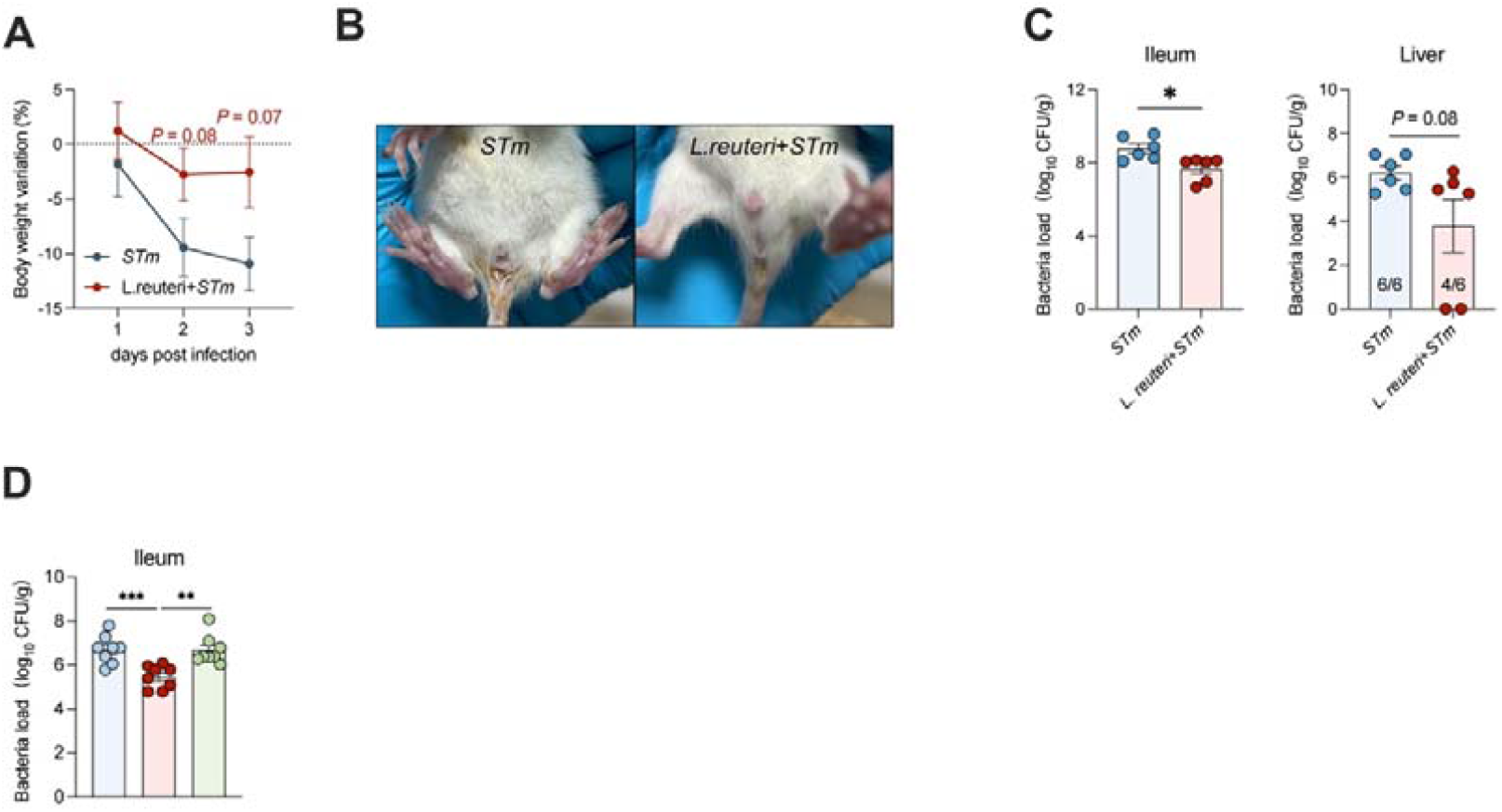
*L. reuteri* protected against *S. typhimurium* infection (A) Bodyweight variation of the *STm* and *L. reuteri* + *STm* groups. The significance of dissimilarity was calculated by t-test. n = 9. (B) Representative diarrhea images of the *STm* and *L. reuteri* + *STm* groups. (C) *STm* burden in the ileum and liver at 3 dpi. The significance of dissimilarity was calculated by t-test. n = 6. (D) *STm* burden in the ileum of *STm*, *L. reuteri* + *STm* and *L. reuteri* + BSH-in + *STm* groups at 1 dpi. The significance of dissimilarity was calculated by t-test. n = 8. Data represent mean ± SEM. \**P*[<[0.05; \*\**P* < 0.01; \*\*\**P* < 0.001. The difference between the two groups was analyzed by two-tailed unpaired Student’s t-test.

## Notes

### Competing Interest Statement

The authors have declared no competing interest.

