## Supplementary Table1 for "Early-life Microbiome-Derived HCA Enhances Type 3 Immunity via FXR- dependent m^6^A RNA methylation"

| **Marker** | **Channel** | **Cat#** | **Company** |
| --- | --- | --- | --- |
| SIRP-α | PE | MA517504 | Thermo |
| CD3 | PE | 550353 | BD Pharmingen |
| CD45RA | PE | 554881 | BD Pharmingen |
| CD127 | Alexa Fluor594 | FAB8484T | R＆D system |
| CD45 | APC-Cy7 | 561586 | BD Pharmingen |
| RORγ (t) | APC | 130-123-840 | Miltenyi |
| IL-22 | PerCP/Cyanine5.5 | 516411 | BD Pharmingen |
| IL-17 | PE-Cy7 | 25-7177-82 | Thermo |
| CD3 | FITC | 559975 | BD Pharmingen |
| CD68 | PE | 130-123-757 | Miltenyi |
| CD45RA | BV421 | 740043 | BD Pharmingen |
| γδ T | BV650 | 745392 | BD Pharmingen |
| CD8 | BV786 | 740913 | BD Pharmingen |
| CD4 | APC | 550057 | BD Pharmingen |
| KLRG1 | AF647 | sc-32755 | Santa Cruz Biotechnology |

Supplementary Table1: Antibody used for flow
