## Supplementary Table2 for "Early-life Microbiome-Derived HCA Enhances Type 3 Immunity via FXR- dependent m^6^A RNA methylation"

**Supplementary Table 2:** Primers used for RT-PCR

| Gene | Primers sequence (5′-3′) | Size (bp) | Accession no. |
| --- | --- | --- | --- |
| β-actin | F: GCAGGAGTACGATGAGTCCG  R: ACGCAGCTCAGTAACAGTCC | 74 | [NM_031144.3](https://www.ncbi.nlm.nih.gov/entrez/viewer.fcgi?db=nucleotide&id=402744873) |
| VEGFA | F: CTCCACCATGCCAAGTGGTC  R: TCACCACTTCATGGGCTTTCT | 70 | NM_001110333.2 |
| SLC2A1 | F: CATCCACCACACTCACCACACT  R: AAACCCATAAGCACGGCAGACA | 166 | NM_138827.2 |
| TFF3 | F: GATAACCCTGCTGCTGGTCCTG  R: CCACGGTTGTTACACTGCTCTG | 150 | NM_013042.2 |
| MUC3 | F: AGTGCTGTTGGTGATTCTTGTG  R: GATGCTCTGCCTTCCTCTTCA | 136 | XM_063271879.1 |
| PFKFB3 | F: CGACAAATGCGACAGGGACTTG  R: ACACGATGCGGCTCTGGATG | 101 | XM_017600444.3 |
| DEFB1 | F: AAGTCTTGGACGCAGAACAGAT  R: AGTTTGGTATGAGATGGGCAGC | 87 | NM_031810.2 |
| BNIP3L | F: AACAACAACTGCGAGGAAGG  R: TGTGGATGGAAGACGAGGAA | 164 | NM_080888.2 |
| HIF-1α | F: CTGACTCTGCTAGCTCCAG  R: TTCAGTTTCCGTGTCATCGC | 75 | NM_024359.2 |
| IL-17A | F: GTGAAGGCAGCGGTACTCAT  R: GGGTGAAGTGGAACGGTTGA | 162 | NM_001106897.1 |
| IL-22 | F: ATACATCGTCAACCGCACCT  R: TGTAAGGCTGGAATCTGTCTGA | 194 | NM_001191988.1 |
| RORγt | F: GCAAAGAAGACCCACACCTCAC  R: GCCGAACTTGACAGCATCTCT | 197 | NM_139089.2 |
| T-bet | F: AATGACGGTGAGCCAGAGG  R: GTAGGCAGTCACGGCAATG | 87 | NM_001107043.1 |
| GATA3 | F: CTCCAGTCCGCATCTCTTCA  R: ACCTGATACTTGAGGCACTCTT | 129 | NM_133293.2 |
| FGF19 | F: GGCTGATTCGCTACTCGGAGGA  R: TGTGGAGGTGGTGCTTCATGGA | 96 | NM_130753.2 |
| SHP | F: AGGAGGCTCACTGGGCATTGT  R: CGATGACAGGGCGGAAGAAGAG | 147 | NM_057133.1 |
| WTAP | F: CATCCTTGTCATGCGGCTAG  R: CACTCGGCTGCTGAACTTG | 92 | XM_063270739.1 |
| METTL3 | F: CCATCCGTCTTGCCATCTCTA  R: CTTGGAGTGGTCAGCATAGGT | 161 | NM_001024794.1 |
| METTL14 | F: TTCGGGAGGGACAGCACTA  R: TCAGACTTGGATTTGGGAGGAG | 182 | NM_001399249.1 |
| ALKBH5 | F: TCGTGTCCGTGTCTTTCTTTAG  R: GTGATCTCATCAGCAGCATACC | 151 | NM_001191643.1 |
| FTO | F: TAAGAGCAGAGCAGCCTACAA  R: TGTCCACCAAGTTCTCGTCAT | 137 | NM_001039713.1 |
| YTHDF1 | F: ATCTGCTCTTCAGTGTCAATGG  R: AGGCTTGTTGTCGTTATTCTCC | 200 | NM_001024756.1 |
| YTHDF2 | F: ACACAGCCATTGCCTCCA  R: TGTCCTACTCCATTACCATCCA | 143 | NM_001047099.2 |
